# Evolution of songbird vocal imitation involved multiple cellular innovations

**DOI:** 10.64898/2026.04.28.721177

**Authors:** Carlos G. Orozco, Ziran Zhao, Ashwinikumar Kulkarni, Judith James, Devin P. Merullo, Genevieve Konopka, Todd F. Roberts

## Abstract

A central goal of evolutionary neuroscience is to link behavioral adaptations to changes across the central nervous system. The dorsal telencephalon (i.e., pallium) varies widely in macrostructure across vertebrate clades and is thought to underlie the diversity of complex vertebrate behaviors. Yet, pallial cell types appear comparatively constrained in their evolutionary specializations relative to other parts of the nervous system, even across species with notable behavioral adaptations. Instead, differences among pallial neurons largely reflect their topographic organization. Here, we show that although transcriptomic identities of cell types in the songbird pallium largely follow this pattern, multiple cellular innovations exist within the vocal imitation brain regions distributed across the pallium. These include behavior-linked specializations to glutamatergic cell types in each region and the presence of PV- and SST-like GABAergic neurons unique to vocal imitation circuits. Additionally, we identify a concerted increase in glial composition that is accounted for by elevated proportions of astrocytes across vocal imitation circuits. These findings demonstrate that complex learned behaviors, such as vocal imitation, can be associated with the emergence of novel, behavior-associated cell types in the pallium; suggesting that selective pressures can, in some uncommon cases, favor functional specialization over conserved developmental constraints.

## Introduction

The evolution of new behaviors can, in some cases, be linked to the emergence of distinct and novel cell types (*1–3*). Evidence for such links has come primarily from peripheral sensory systems (*4–10*), the invertebrate central nervous system (*11–13*), and even from modulation from outside the nervous system (*14*). However, it remains uncertain whether cognitively complex behaviors can be directly linked to cellular innovations within higher-order brain regions or if these circuits are inherently more constrained.

One illustrative example is the pallium, a major forebrain division shared across vertebrates that integrates sensory inputs, supports motor control, and underlies complex cognition. Across amniotes (mammals, birds, and reptiles), disproportionate pallial expansion largely accounts for differences in brain size and is associated with behavioral repertoires (e.g., the human neocortex). However, identified clade-specific cellular innovations are remarkably conserved within clades and, to our knowledge, lack clear links to species-specific complex behaviors. For example, chandelier cells appear to be mammalian-specific (*15*), Rosehip interneurons have so far only been found in humans (*16*), and the mammalian expansion of upper-layer intratelencephalic neurons correlates with increasing cognitive complexity (*17*). Yet, mammalian pallial cell-type heterogeneity has been correlated to spatial topography rather than to specific behaviors (*15–18*). Thus, developmental constraints in the pallium may favor conservative modification of general-purpose cell types rather than the emergence of new ones.

Examining evolutionarily rare and specialized behaviors provides an opportunity to directly test if they can be linked to specific cellular innovations. Songbirds present a model for testing whether complex behavioral repertoires evolve in association with specific cellular specializations. The songbird brain contains an interconnected set of pallial regions important for learning and producing song (*19–21*), and are sexually dimorphic in species in which only males learn song, such as zebra finches (*22–26*). These regions are embedded within anatomical areas dedicated to nonvocal but broadly similar functions that form parallel pathways with vocal circuits (*27–35*). This capacity to modify vocalizations through imitation—a prerequisite for both speech and song learning—is rare and has evolved in only a few vertebrate lineages (*36*). Given this phylogenetic rarity and tractability of the songbird brain, this circuitry provides a powerful model for testing whether behavioral novelty is accompanied by the emergence of novel cell types in the higher brain centers of vertebrates. Or, instead whether cell-type heterogeneity is tied to spatial topography.

Previous bulk RNA-seq, in situ hybridization, and related approaches have begun to uncover molecular differences between vocal and nonvocal pallial structures (*37–39*). However, it is unclear if these gene expression differences occur across all cell types in vocal regions, are cell-type specific, or reflect differences in cell-type proportions. Building on these efforts, our previous work used single-nucleus RNA sequencing (snRNAseq) of the premotor and prefrontal cortex analog HVC (proper name) and the vocal-motor cortex analog, RA (robust nucleus of the arcopallium), and revealed distinct cell types analogous to those found in the mammalian cortex (*40*). It remains unclear whether these subtypes are unique to HVC and RA or are found more broadly throughout the brain, as counterpart comparisons were lacking. Indeed, a recent study that compared RA to the rest of the motor cortex analog at single-cell resolution identified molecular specializations of glutamatergic neurons (*41*), highlighting the importance of within-species comparative approaches.

As with many complex behaviors, vocal imitation engages additional brain regions upstream of the motor cortex. To explore whether this behavioral expansion involved the emergence of novel cell types—or instead reflects minimally distinct, regionally distributed cell types—we used snRNA-seq and spatial transcriptomics (sptRNA-seq) (**Supplementary Table 1**). The regions examined encompassed motor and premotor cortices (including prefrontal cortex (PFC) analogs), upstream secondary auditory areas, and a primary visual cortex analog. To test if any differing cell types are indeed evolutionarily novel, we compared cell types between these regions and to cell types in homologous brain regions of chickens, which produce only innate vocalizations and are evolutionarily separated from songbirds by approximately 100 million years (*42*).

We observed innovations in glutamatergic and GABAergic neurons and proportional changes to non-neuronal cells in premotor and motor regions necessary for vocal imitation. In contrast, general motor, auditory, and visual regions, which are conserved across songbirds and chickens, exhibited far less transcriptomic specializations. Together, these results provide the first evidence, to our knowledge, directly linking a complex learned behavior to the emergence of novel, behavior-specific cell types across the vertebrate pallium.

### Transcriptomic profiling of vocal imitation and adjacent brain regions

To identify potential cell types associated with vocal imitation, we used snRNA-seq on multiple pallial regions of six adult zebra finches (133-185 days post-hatch) with mature imitated song (**Supplementary Table 1**). These included three brain regions that have been shown to be essential for imitation and the production of birdsong, and are absent in non-vocal-learning birds: the vocal-motor cortex RA, and the PFC/premotor cortex analog HVC, another premotor region essential for song acquisition, LMAN (lateral magnocellular nucleus of the anterior nidopallium), along with the incorporation of our previously generated dataset from two of these regions (*40*) (**Extended Data Fig. 1A-B**).

To disentangle vocal imitation specializations from possible topographic patterning associated with developmental origin or connectivity, we also sequenced regions with the same developmental origin and parallel connectivity patterns, but that are not involved in vocal imitation. These included: motor cortex analog, AA (anterior arcopallium), to compare with RA; premotor cortex analog, PFC-analog, NCL (nidopallium caudolateral), to compare with HVC; and AN (anterior nidopallium), to compare with LMAN (**Extended Data Fig. 1A**). Distinct cellular profiles in vocal imitation regions may also reflect more extensive specialization in the songbird pallium, or more general adaptations related to sensory processing. To distinguish these possibilities, we included auditory and visual areas present in both songbirds and non-vocal-learning species: specifically, the auditory region NCM (caudomedial nidopallium) to compare with the developmentally related regions NCL and HVC; the auditory region CM (caudal mesopallium) and a developmental counterpart from the AM (anterior mesopallium); and a region of the hyperpallium (hyper) that contains a primary visual cortex analog (**Extended Data Fig. 1A**).

After collecting nuclei from these regions and carrying out snRNA-seq, we removed ambient RNA, doublets, low-nuclear fraction reads, low-quality cells, and sample-specific clusters (**Extended Data Fig. 1C-D**). This resulted in 380,001 nuclei with a median of 1,458 RNA transcripts [unique molecular identifiers (UMIs)] and 935 transcribed genes detected per cell (**Extended Data Fig. 1E-I**). We also generated a complementary sptRNA-seq dataset from seven sagittal sections (n = 3 zebra finches) that included all the pallial vocal imitation regions and most of the controls from our snRNA-seq dataset (**Fig. 1A**). This was used to examine the spatial distributions of snRNA-seq–defined cell types and test if similar cell types are found in regions not captured by the single-nucleus dataset.

**Fig. 1.**
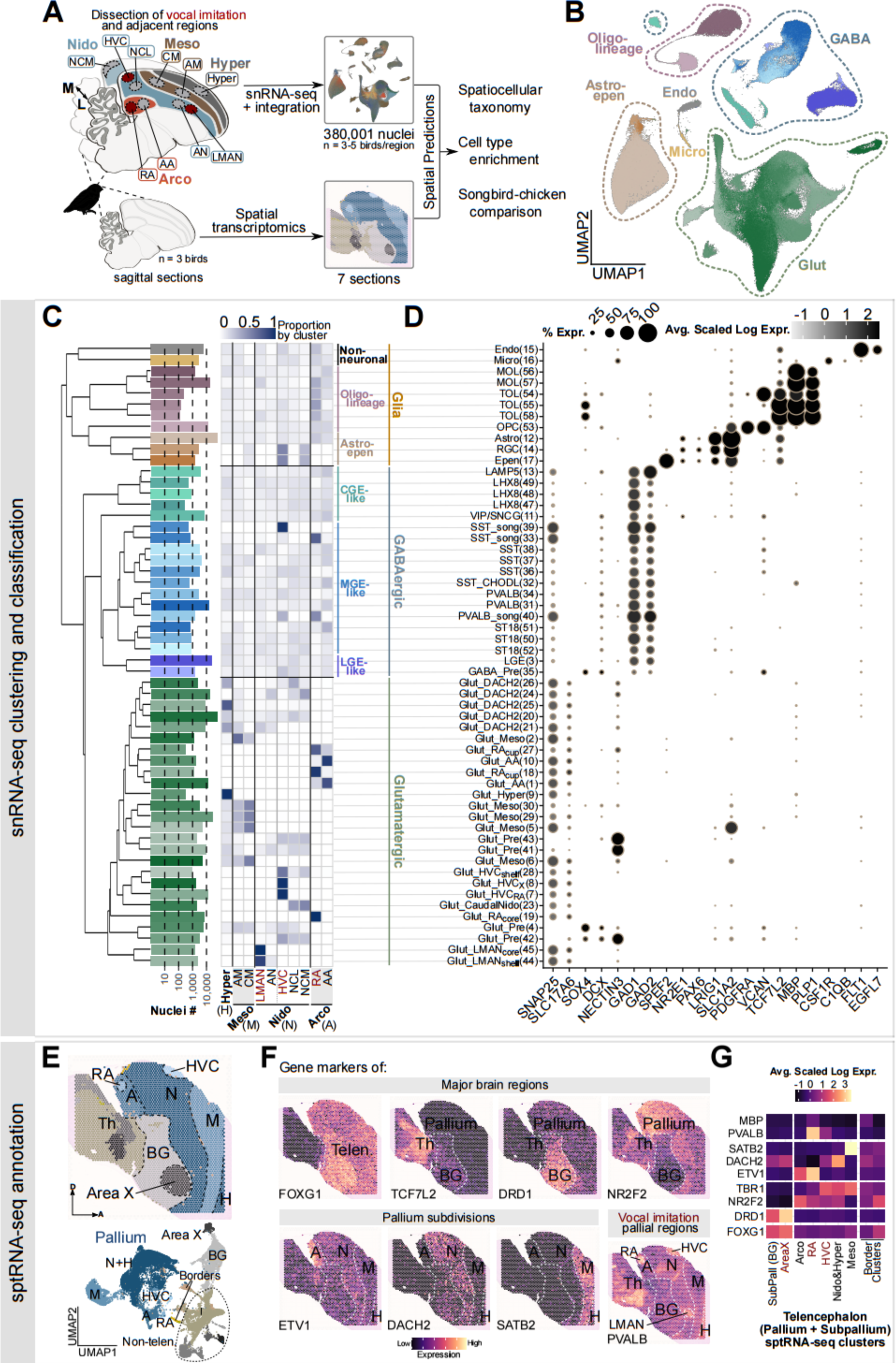
Cellular and spatial transcriptomics of vocal imitation and neighboring pallial areas. (A) Schema of datasets and analysis. Pallial regions are color-coded, and dissections within those regions are enclosed in matching-colored boxes Hyper, hyperpallium; meso, mesopallium; nido, nidopallium; arco, arcopallium; M, medial, L, lateral. (B) Uniform manifold approximation and projection (UMAP) of zebra finch pallium single-nucleus RNA sequencing (snRNA-seq) atlas, colored by cluster annotation as shown in (C). Astro, astrocyte; Epen, ependymal; Glut, glutamatergic neurons; GABA, GABAergic neurons; Oligo, oligodendrocyte; Micro, microglia; Endo, endothelial. (n = 380,001 nuclei, 3-5 birds per region) (C) Left, transcriptomic taxonomy tree of 56 clusters organized in a dendrogram, with attributes from left to right: number of cells colored as in (B); heatmap representing the proportion of cells per cluster from each brain region; cell class and subclass name. (D) Cluster names and gene expression dotplot of genes that identify neurons (*SNAP25*); glutamatergic (*SLC17A6*), immature (*SOX4*, *DCX*), and GABAergic (*GAD1*, *GAD2*) neurons; ependymal cells (*SPEF2*, *PAX6*); astrocytes (*LRIG1*, *SLC1A2*); OPCs (*PDGFRA*, *VCAN*); transitioning (*FYN*) and mature oligodendrocytes (*MBP*, *PLP1*); immune cells (*C1QB*, *CSF1R*); and endothelial cells (*FLT1*, *EGFL7*). Astro, astrocytes; MOL, mature oligodendrocytes; TOL, transitioning oligodendrocytes; OPC, oligodendrocyte precursor cells. Expression less than 5 percent is excluded for clarity. (E) Top, spatial plot displaying clustering of spots of an example sagittal section. Bottom, integrated UMAP of all zebra finch sagittal brain spatial RNA sequencing sections. A, arcopallium; N, nidopallium; M, mesopallium; H, hyperpallium; BG, basal ganglia; Telen, telencephalon; Th, thalamus. (F) Spatial expression (log-normalized) plots of genes *FOXG1*, *TCF7L2*, *DRD1*, *NR2F2*, *ETV1*, *DACH2*, *SATB2*, *PVALB*, used to mark the telencephalon, thalamus, basal ganglia, pallium, arcopallium, nido- and hyper-pallium, mesopallium, and song regions, respectively, in an example sagittal section. Regional borders are marked by a white outline. (G) Heatmap of the average scaled expression (log-normalized) across all sagittal sections by spatial cluster of marker genes (except *TCF7L2*) shown in (F), plus *TBR1*, another pallium gene marker, and *MBP*, a myelin gene marker. SubPall, subpallium.

Zebra finch snRNA-seq data were clustered using an iterative algorithm based on differential gene expression thresholds, and cluster stability was assessed through repeated bootstrapping (*43, 44*), yielding 56 stable clusters. We then annotated them based on canonical marker genes and grouped them into six major cell classes: glutamatergic neurons, GABAergic neurons, astroependymal cells, oligodendrocyte lineage cells, microglia, and endothelial cells (**Fig. 1B-D**). This included 1 immature GABAergic neuron subtype found in all regions, GABA_Pre(35). It also included 4 immature glutamatergic neuron subtypes (Glut_Pre(4 & 41-43)) found in all dissection regions except the arcopallium, consistent with previous reports of robust neurogenesis in avians in some but not all pallial regions (*45–47*). Additionally, there were 22 subtypes of mature glutamatergic and 18 GABAergic neurons (**Fig. 1C-D**).

To obtain an anatomically relevant delineation of our sptRNA-seq data (**Supplementary Table 1**), we used low-resolution clustering. Together with anatomical location and marker gene expression, we delineated major brain regions: the thalamus (*TCF7L2*); the telencephalon (*FOXG1*) and its ventral subregion, subpallium or basal ganglia (*DRD1*); and its dorsal subregion, the pallium (*NR2F2* and *TBR1*). Within the pallium, we then identified the arcopallium (*ETV1*), nidopallium and hyperpallium (*DACH2*), mesopallium (*SATB2*), and the vocal-imitation nuclei HVC, LMAN, and RA (*PVALB*) (**Fig. 1E-G** and **Extended Data Fig. 2**).

### Vocal-imitation glutamatergic neurons have unique transcriptomic signatures

Our sptRNA-seq demonstrated that even at a coarse scale, HVC and RA emerged as discrete clusters, distinct from the surrounding nidopallium and arcopallium, respectively (**Fig. 1E-G**). This segregation in high-dimensional transcriptomic space (30 principal components) underscores their pronounced gene expression differences, highlighted by their relatively small size and the absence of other separate clusters within the larger pallial regions in which they are embedded. Since these two regions belong to the vocal motor pathway, this raises the possibility that motor control of vocalizations represents one of the most specialized functions within the pallium. Additionally, given the relatively coarse resolution of sptRNA-seq and the high density of glutamatergic neurons in these regions, glutamatergic neurons are likely the main contributors to the observed transcriptomic divergence.

Supporting this, our snRNAseq data show that half of the glutamatergic subtypes (11/22) were strongly biased toward a single dissection region (≥70% of cells from one dissection region). Of these, the majority (7/11) were from the brain regions that form part of the vocal imitation regions: RA, HVC, or LMAN (**Fig. 1C-D**). This indicates that glutamatergic neurons in vocal imitation regions exhibit stronger categorical, region-specific transcriptomic specialization relative to other glutamatergic neurons in our dataset, whose regional differentiation might be more strongly associated with anatomical topography rather than behavior (*15–18*).

A continuous measure, agnostic to clustering resolution, also revealed stronger regional transcriptomic signatures of vocal imitation glutamatergic neurons. We quantified the degree of mixing in the transcriptomic space between each dissection region by computing the Local Inverse Simpson’s Index (LISI)—which scores each cell, with higher values indicating greater mixing and thus higher gene-expression similarity (*48*). We grouped cells into major cell classes and into those originating from vocal imitation regions (HVC, LMAN, or RA) or nonvocal regions. In both groupings, glutamatergic neurons had the lowest median LISI score among the major cell classes (excluding ependymal cells)—indicating that they are transcriptomically more distinct across regions than other major cell classes, a pattern consistent with previous reports (*15, 17, 18, 40, 49*). However, glutamatergic neurons of vocal imitation regions had lower LISI scores than nonvocal regions (median of 1.7 vs. 2.6) (**Fig. 2A-B**). These results further support increased regional specialization of glutamatergic neurons in vocal imitation regions, because lower LISI scores indicate less mixing in transcriptomic space across regions and therefore more distinct, region-specific transcriptional profiles.

**Fig. 2.**
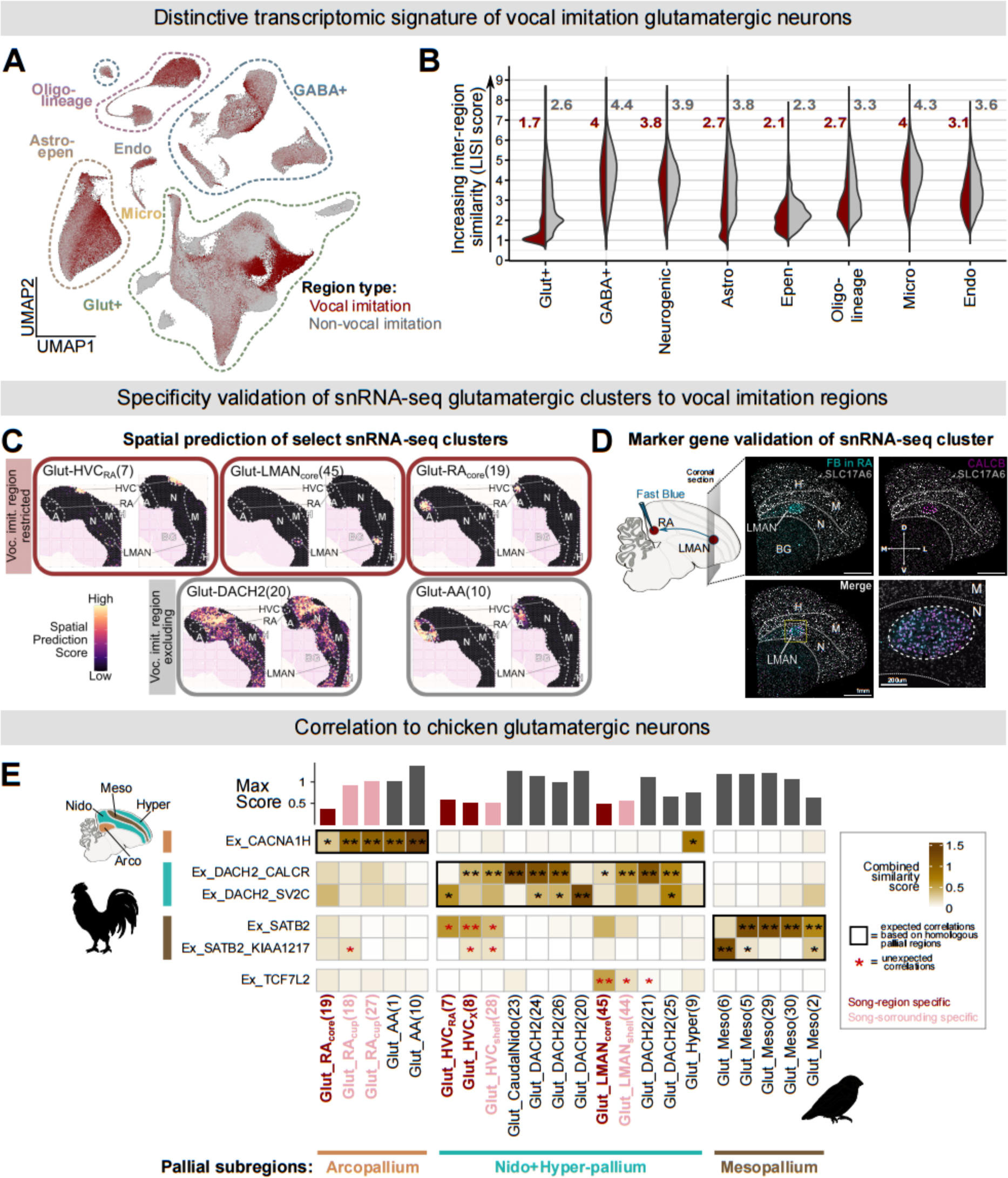
Specialization and divergence of vocal-imitation glutamatergic neurons. (**A**) UMAP representation of snRNA-seq cell types colored by song region (red) or other (grey). Outlines grouping them by major cell classes. (**B**) Violin plots of local inverse Simpson’s index (LISI) indicating the degree of mixing in 30 principal component transcriptomic space between cell classes, coming from either song (red) or other (grey) regions, where higher scores reflect greater mixing (higher transcriptomic similarity) and lower scores indicate minimal mixing (lower transcriptomic similarity). Numbers indicate median LISI score. (**C**) Spatial prediction scores of example clusters predominantly composed of cells from either song regions (red outline) or control regions (grey outline). High spatial prediction scores indicate a high probability that cells with the respective identity were present within the spot’s area. (**D**) Schematic of retrograde tracing analysis via injection of fast blue into the LMAN target RA. Split channels from an example coronal section of a bird injected with Fast Blue in RA or Glut_LMANcore(45) neurons’ marker (*CALCB*), combined with in situ hybridization for glutamatergic neuron marker (*SLC17A6*), and merged image. (**E**) Comparison of chicken glutamatergic subclasses and zebra finch clusters using combined scores from CCA integration label transfer and Spearman correlation of gene specificity index for differentially expressed genes. Bar plots at the top show the highest similarity scores. Black outline boxes represent expected correlations based on the homology of pallial regions.

Spatial mapping using Canonical Correlation Analysis (CCA) integration with sptRNA-seq broadened support for regional specificity, showing that these signatures did not generalize to unsampled regions by snRNA-seq. Namely, the most restricted spatial mapping (prediction scores) was from ‘core’ song-region clusters—Glut_LMAN_core_(45), Glut_RA_core_(19), Glut_HVC_RA_(7), and Glut_HVC_X_(8). This was similar in ‘shell’ clusters from co-dissected regions immediately adjacent to vocal imitation regions (see Technical Note in Methods)—Glut_HVC_shelf_(28), Glut_LMAN_shell_(44), Glut_RA_cup_(18), and Glut_RA_cup_(27). In situ hybridization for cluster-specific marker genes further corroborated this spatially restricted distribution (**Fig. 2C-D** and **Extended Data Fig. 3A-G**).

In contrast, glutamatergic clusters from outside vocal-imitation regions showed spatial mapping that reflected developmental relationships. For example, Glut_DACH2(20) and Glut_AA(10) mapped broadly across their respective pallial regions, the nidopallium and arcopallium. Similarly, clusters with many cells from the auditory regions NCM and CM (**Fig. 1C**) mapped to their corresponding pallial regions, namely the nidopallium and mesopallium, respectively (**Fig. 2C**). This suggests that the primary transcriptomic signature of pallial glutamatergic neurons outside song regions reflects their developmental origin, akin to the topographic patterning of glutamatergic neurons in mammals. These findings contrast with the singular transcriptomic profiles of glutamatergic neurons in vocal imitation areas; further suggesting that these neurons may reflect behavior-associated cellular innovations that emerged during the evolution of vocal imitation.

### Divergence of vocal imitation glutamatergic neurons from ancestral cell types

To better examine if these cell types are indeed novel and unique to vocal learning brain regions, we compared them to a published snRNA-seq dataset of the chicken pallium (*Gallus gallus* (*49*)), an avian species that lacks vocal imitation brain regions but has developmentally homologous pallial subdivisions. We compared their glutamatergic neurons using label transfer with CCA (*50*) and gene specificity index (GSI) correlations (*51*), and aggregated them into a single score. To increase confidence and robustness of the correlations, we mainly focused on the coarser annotation level of the chicken dataset (**Fig. 2E** and **Extended Data Fig. 4A-B**).

We found that the songbird clusters derived from non-vocal-imitation regions had stronger correlations to chicken neuronal subclasses and they were mainly associated with homologous brain regions. For example, the auditory region, CM, formed five mixed clusters, Glut_Meso(6, 5, 29, 30, and 2), with the developmentally related AM region within the mesopallium. These zebra finch clusters only had significant correlations to the identified chicken subclasses, Ex_SATB2 or Ex_SATB2_KIAA1217, which were localized to the chicken mesopallium (**Fig. 2E** and **Extended Data Fig. 4A-B**), suggesting the mesopallium clusters are mainly conserved across species.

Moreover, clusters Glut_DACH2(26, 24, 25, 20, 21, and 23) contained cells primarily originating from dissections within the nidopallium or hyperpallium regions, consistent with the known molecular similarity between these regions (*49, 52*), and thus not segregated based on their functional specialization. These clusters had high correlations to the chicken nidopallium/hyperpallium clusters Ex_DACH2_CALCR and Ex_DACH2_SVC2 (**Fig. 2E**). Notably, the small cluster Glut_Hyper(9) showed significant correlation only with the chicken arcopallium subclass, despite being specific to the hyperpallium. This apparent discrepancy was clarified at higher resolution: the zebra finch cluster strongly correlated with the chicken hyperpallium-specific subtype Ex_TSHZ2_NR4A2, a small cluster with gene-expression similarities to arcopallial glutamatergic neurons that mapped to outward-projecting neurons of the apical hyperpallium (**Extended Data Fig. 5A-C**).

By contrast, nidopallium clusters stemming from the vocal imitation premotor region LMAN, Glut_LMAN(44 & 45), had lower maximum scores to chicken nidopallium glutamatergic clusters. Instead, Glut_LMAN_core_(45) had the strongest correlation to subclass Ex_TCF7L2 (**Fig. 2E** and **Extended Data Fig. 4A-B**). This chicken neuron subclass was described as likely arising from the thalamus due to an inadvertent dissection mistake; evidenced by transcription factor expression of *TCF7L2*(*53*), small cluster size, and dendrogram-based gene expression similarity to GABAergic neurons (*49*). Yet, our correlations suggest that Ex_TCF7L2 may represent a rare avian cell type that was evolutionarily co-opted and expanded to become LMAN, which functions as the cortical output pathway of the basal ganglia loop essential for vocal imitation (*54–58*).

Clusters from the other nidopallium vocal imitation region, HVC, had weaker correlations to the chicken cell types. Namely, the basal ganglia-projecting Glut_HVC_X_(8), motor-projecting Glut_HVC_RA_(7), and the Glut_HVCshelf(28) clusters had weaker correlations to the homologous chicken brain region and had significant correlation to either the Ex_SATB2 or Ex_SATB2_KIAA1217 subclasses that are restricted to the mesopallium, not the nidopallium (**Fig. 2E** and **Extended Data Fig. 4A-B**). These correlations suggest an exaptation and molecular module integration (*2, 59*), in which a core regulatory complex(es) in ancestral mesopallium cell types may have been co-opted by the HVC projection neurons to produce novel cell types.

In addition, the Glut_RA_core_(19) cluster, representing the projection neurons of the vocal motor cortex, displayed correlations distinct from those of nonvocal motor cortex clusters—Glut_Arco(1), Glut_RA_cup_(18), Glut_AA(10), and Glut_RA_cup_(27)—despite all residing in the arcopallium. Specifically, Glut_RA_core_(19) had the lowest maximum prediction score from all the arcopallium clusters. Additionally, one of the RA shell (or cup) clusters, Glut_RA_cup_(18), had a significant correlation to the chicken mesopallium subclass. In contrast, clusters from the zebra finch arcopallium outside of RA were highly correlated with the chicken arcopallium subclass Ex_CACNA1H (**Fig. 2E** and **Extended Data Fig. 4A-B**). Therefore, the vocal motor cortex appears to have diverged through a mechanism distinct from the co-option proposed for LMAN and HVC cell types.

Overall, this analysis reveals that vocal-imitation-related glutamatergic subtypes show weak cross-species correlations, pointing to behavior-specific innovations. In contrast, glutamatergic subtypes from auditory, visual, and general motor regions in zebra finches exhibited transcriptomic similarity to cell types in homologous pallial regions of chickens. Additionally, the potential mechanisms of these vocal imitation cellular innovations might have been unique to each region.

### Unique divergence and distinct evolutionary mechanisms of glutamatergic neurons

Indeed, the LISI scores grouped cluster were more suggestive of a unique transcriptomic signature rather than a shared song-related gene regulatory program. Song clusters tended to have scores closer to 1 than their counterpart non-song clusters, thus supporting regional transcriptomic uniqueness, even compared to other song regions (**Fig. 3A**). A notable contrast was the cluster with the most neurons, Glut_DACH2(20), which had a similar percentage of nuclei originating from all non-arcopallium dissections (**Fig. 3A, Extended Data Fig. 6A**, and **Supplementary Table 2**). This cluster mainly consisted of neurons previously identified as non-projecting and putatively immature, based on the absence of known projections in our earlier study (*40*) (**Extended Data Fig. 1B**). This indicates that immature or locally projecting glutamatergic neurons might be among the most prevalent neuron types in the pallium with minimal transcriptomic distinction between regions. Although it is unclear whether they are immature non-projecting (*60–63*) or solely locally projecting (*63–65*).

**Fig. 3.**
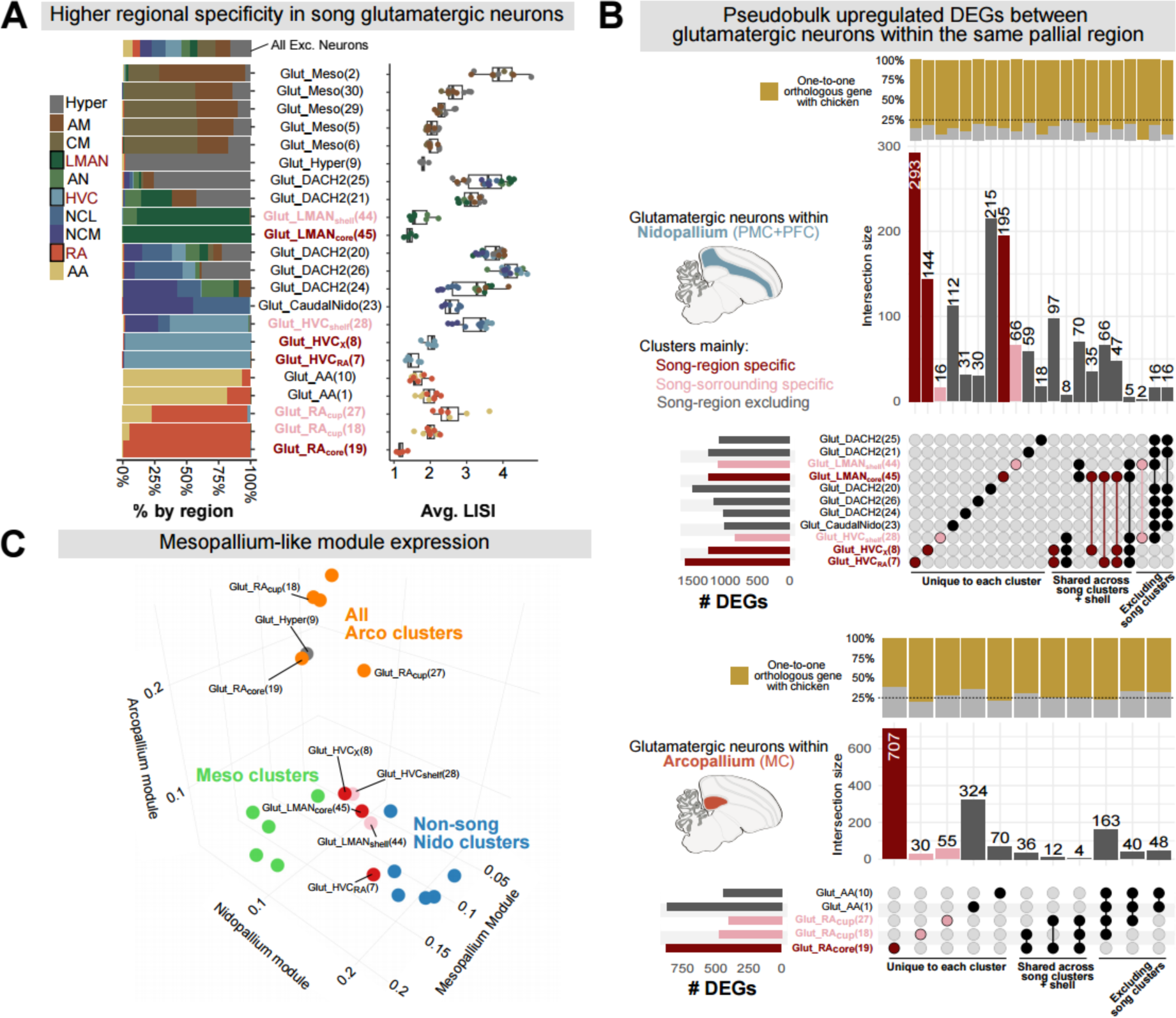
Regional specialization of glutamatergic neurons in vocal imitation areas. (**A**) Left, stack plots showing the fraction of cells profiled by region per cluster and all glutamatergic neurons. Right, LISI scores by replicate, including only replicates that contributed more than five percent of the cells in a given cluster. (**B**) Top, upset plot of pseudobulk differentially expressed genes for nidopallium clusters, defined by pairwise comparisons of each cluster against all others within the region. The top vertical bars show the percentage of genes with 1-to-1 orthologs in the chicken genome in the corresponding intersection. Bottom, the same analysis for arcopallium clusters. PMC, premotor cortex analog; PFC, prefrontal cortex analog. MC, motor cortex analog (C) Comparison of nidopallium, mesopallium, and arcopallium module scores for all glutamatergic neuron clusters.

We further distinguished between distinct or shared gene regulatory programs through pseudobulk differential gene expression comparisons. Song-restricted glutamatergic neurons from vocal imitation regions exhibited the highest number of differentially expressed genes (DEGs), with each cluster also showing the greatest number of unique DEGs relative to other clusters (**Fig. 3B, Extended Data Fig. 6B** and **D**, and **Supplementary Table 3**). The predominance of unique over shared DEGs reinforces our previous inference that each of these novel neurons underwent distinct modes of divergence, rather than a distinctly shared regulatory shift. Notably, among the few shared upregulated DEGs was the sexually dimorphic transcription factor *AR*, which was differentially expressed in LMAN projection neurons and in both major HVC projection neuron types. This shared set also included autism risk genes such as *ASTN2* (*66*) and *DPP6* (*67*), consistent with the involvement of these circuits in vocal learning, a domain frequently disrupted in autism spectrum disorders (ASD) (*36, 68*) (**Supplementary Table 3**).

Moreover, while most differentially expressed genes had 1-to-1 orthologs with chicken genes, those in the arcopallium clusters had a higher percentage of non-orthologous genes (**Fig. 3B** and **Extended Data Fig. 6B-C**). This, together with the lower similarity scores to the chicken arcopallium, suggests that the song motor cortex analog may have specialized not only through regulatory changes but also through coding-sequence innovation. The same cross-species comparisons suggested that vocal imitation clusters in the nidopallium exhibited gene expression patterns like those of unrelated developmental regions, such as the mesopallium. To further test this, we defined gene modules for glutamatergic neurons in each pallial region using pseudobulk pallial-region comparisons that excluded song regions (**Extended Data Fig. 6D** and **Supplementary Table 4**). Song clusters within the nidopallium, indeed, had higher mesopallium module scores than the rest of the nidopallium glutamatergic clusters (**Fig. 3C**) and further supported the possible co-option of a cross-region expression module.

Taken together, these findings suggest that the nidopallium may possess an unusual degree of flexibility in its cellular evolutionary mechanisms. Previous work in chickens showed that gene expression convergence between the nidopallium and hyperpallium occurs throughout development (*49*). Here, we extend this concept by demonstrating that glutamatergic song neurons also converge on gene expression programs characteristic of the mesopallium, a region associated with higher cognitive functions (*69*). These results suggest that nidopallial vocal imitation neurons may have largely emerged through co-option of regulatory programs from developmentally or functionally distinct regions, as well as from rare cell types such as Ex_TCF7L2. In contrast, the arcopallial motor cortex analog appears to have diverged primarily through de novo mechanisms and using expression programs not present in the most recent common ancestor of songbirds and chickens. Thus, highlighting that distinct evolutionary strategies can give rise to specialized neural circuits supporting complex learned behaviors.

### Specialized MGE-like GABAergic neurons in vocal imitation regions

Unlike glutamatergic neurons, most GABAergic neuron clusters were broadly distributed across all dissection regions, consistent with previous cross-species studies (*18, 40, 49, 51*). Unexpectedly, three of the eighteen GABAergic subtypes were strongly biased to specific dissection regions, with most of their cells originating from one or two vocal-imitation regions (**Fig. 4A, Extended Data Fig. 7A**, and **Supplementary Table 5**). LISI scores for these GABAergic neurons were lower than the other clusters (**Fig. 4A**). To quantify regional enrichment, we calculated each cluster’s fraction of GABAergic neurons per region and compared it to the global average and to their paired nonvocal regions. All three subtypes showed markedly higher representation in vocal premotor and motor areas (**Fig. 4B** and **Supplementary Table 5**). This specialization is unlikely to arise from a simple expansion of GABAergic neuron number, as these regions instead exhibited a lower or similar GABAergic-to-glutamatergic neuron ratio relative to their nonvocal counterparts (**Extended Data Fig. 7B** and **Supplementary Table 5**).

**Fig. 4.**
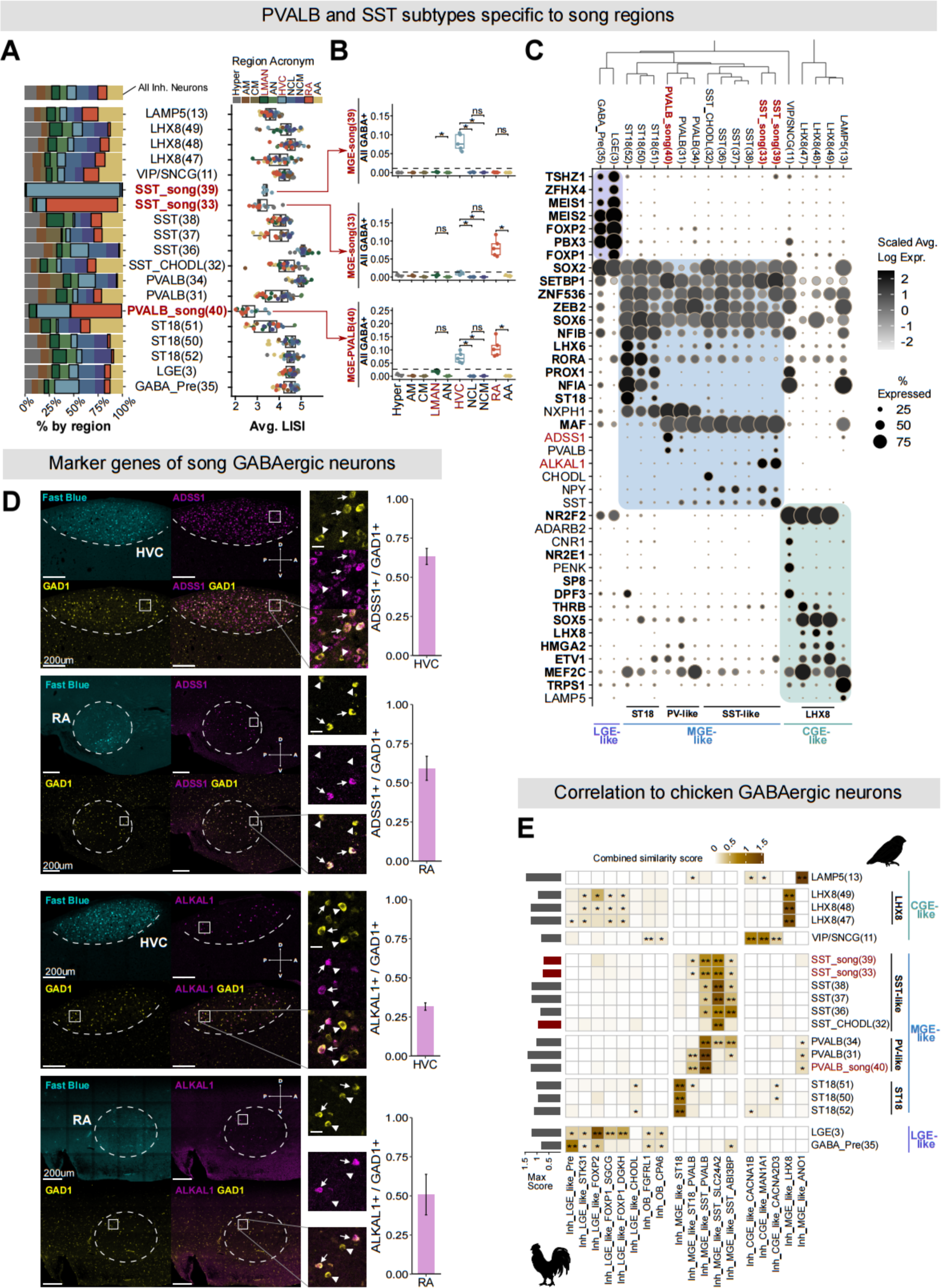
GABAergic neurons of the songbird pallium and specialized song MGE-derived subtypes. (**A**) Left, stack plots showing the fraction of cells profiled by region per cluster and all GABAergic neurons, with song regions outlined in black. Right, LISI scores by replicate, including only replicates that contributed more than five percent of the cells in each cluster. (**B**) Ratio of GABAergic to glutamatergic neurons shown by region, with Wilcoxon rank-sum tests comparing song regions to their anatomical counterparts. Dashed line indicates the average ratio across all regions. (**C**) Dendrogram of GABAergic cell clusters, with dot plot below of marker genes for known GABAergic subclasses. Transcription factors are bold, and genes validated in (D) are shown in red. Less than 1 percent expression is excluded for clarity. (**D**) Example sagittal sections of retrogradely identified HVC or RA (outlined), combined with *in situ* hybridization of GABAergic neuron marker, *GAD1*, and song-specific GABAergic subtype marker genes, *ADSS1* (PVALB_song(40)) and *ALKAL1* (SST_song(33 & 39)). Scale bar, 200 μm. Boxed regions are enlarged at right. Scale bar, 10 μm. Arrows in enlargements indicate examples of *GAD1+* and marker gene-positive cells, while arrowheads indicate *GAD1+* but marker gene-negative cells. Right, quantification of marker gene presence in *GAD1+* cells within the retrogradely identified region. (**E**) Comparison of chicken GABAergic subclasses and zebra finch subtypes using combined scores from CCA integration label transfer and Spearman correlation of gene specificity index for differentially expressed genes. Bar plots on the left show the highest prediction scores. Color bars on the right represent the putative developmental origin region based on marker genes.

Since major GABAergic subclasses are broadly conserved across vertebrates, we next asked to which subclasses these vocal-imitation enriched types belonged. Using canonical markers and differentially expressed transcription factors, we identified the expected GABAergic subclasses derived from the lateral, medial, and caudal ganglionic eminences (LGE, MGE, and CGE), and their corresponding subclasses PVALB-like, SST-like, ST18, LHX8-like, and LAMP5-like GABAergic neurons (**Fig. 4C**). MGE-derived GABAergic neurons include parvalbumin- and somatostatin-GABAergic neurons and are generally conserved across large evolutionary distances. In both our dataset and the chicken dataset (*49*), the MGE lineage also included an ST18-expressing subclass that formed a distinct branch within the MGE-derived populations. Marker genes for this group included the transcription factors NFIB, RORA, and LHX6, which are also upregulated in chandelier cells (*70, 71*) (**Fig. 4C**).

The taxonomy, canonical markers, and a heatmap of differentially expressed genes consistently assigned the enriched GABAergic neurons to either the PVALB-like subclass (PVALB_song(40)) or the SST-like subclass (SST_song(33) and SST_song(39)) (**Fig. 4C, Extended Data Fig. 7C**, and **Supplementary Table 6**). We identified genes *ADSS1* as a marker for PVALB_song(40) and *ALKAL1* for SST_song(33) and SST_song(39) (**Fig. 4C**). We independently validated these vocal imitation–restricted GABAergic neuron subtypes using fluorescence in situ hybridization for these marker genes. Consistent with the snRNA-seq data, expression of *ADSS1* and *ALKAL1* was confined to HVC and RA and observed in only a subset of GAD1⁺ cells (**Fig. 4D**). Notably, these same marker genes were reported in a previous snRNA-seq study of HVC, which identified one of the SST subtypes expressing *ALKAL1* and the only PV subtype expressing *ADSS1* (*72*). Our inclusion of additional pallial regions enabled us to highlight that the SST⁺ALKAL1⁺ and PV⁺ADSS1⁺ populations are vocal imitation–specific GABAergic neuron subtypes.

We next compared zebra finch and chicken pallial GABAergic neurons using the same cross-species correlation framework we applied to glutamatergic neurons. At a broad level, song-related GABAergic neurons aligned with the MGE subclass, like their sister subtypes. At a finer level, PVALB_song(40) most strongly matched the chicken Inh_MGE_like_SST_PVALB cluster, which in turn aligned with mammalian PVALB GABAergic neurons. Conversely, SST_song(33) and SST_song(39) corresponded to the Inh_MGE_like_SST_SLC24A2 chicken cluster with the strongest similarity to mammalian SST GABAergic neurons (**Fig. 4E** and **Extended Data Fig. 7D-E**).

These results show that, alongside with the widespread GABAergic cell types, vocal-imitation regions contain evolutionarily novel MGE-derived subtypes with distinctive transcriptomic features. Furthermore, the presence of identifiable sister cell types across regions and species underscores the conservation of GABAergic neurons, while also demonstrating that behavioral innovation may arise through transcriptomic changes even in deeply conserved cell types.

### Transcriptomic specialization of song region-specific neurons

The GABAergic clusters enriched in vocal imitation regions formed part of the SST- and PVALB-like lineages (**Fig. 4C**). We therefore focused on lineage-specific expression changes by performing pseudobulk differential gene expression analysis (DESeq2, logFC > 0.1, adj. p < 0.05) and comparing song-region PV- and SST-like GABAergic neurons to their more widespread PV-and SST-like counterparts. This showed that many of the upregulated genes in song neurons had 1-to-1 orthologs in the chicken genome (74% across all comparisons, except in PV vs PV-song, in which it was 77%), indicating that the programs to differentiate from other similar subtypes were largely driven by changes in gene expression rather than in coding sequences.

We examined transcription factors, which regulate gene expression and thus cellular identity (*18, 59*), and on autism-risk genes, given that disruptions in vocal communication are central features of ASD (*36, 68*). This analysis revealed only six transcription factors upregulated in song-related GABAergic neuron types, including *CUX1* and *SETBP1*, which have been associated with autism risk (SFARI). Additionally, *CUX1* has been shown to be regulated by a human accelerated region and demonstrated to play a role in increasing synaptic spine density and area in an activity-dependent manner (*73*). We also identified *RUNX1*, a transcription factor implicated in the neuronal diversification of dorsal root ganglion neurons, where it regulates fine laminar termination and suppresses CGRP expression (*74*) (**Fig. 5A** and **Supplementary Table 7**).

**Fig. 5.**
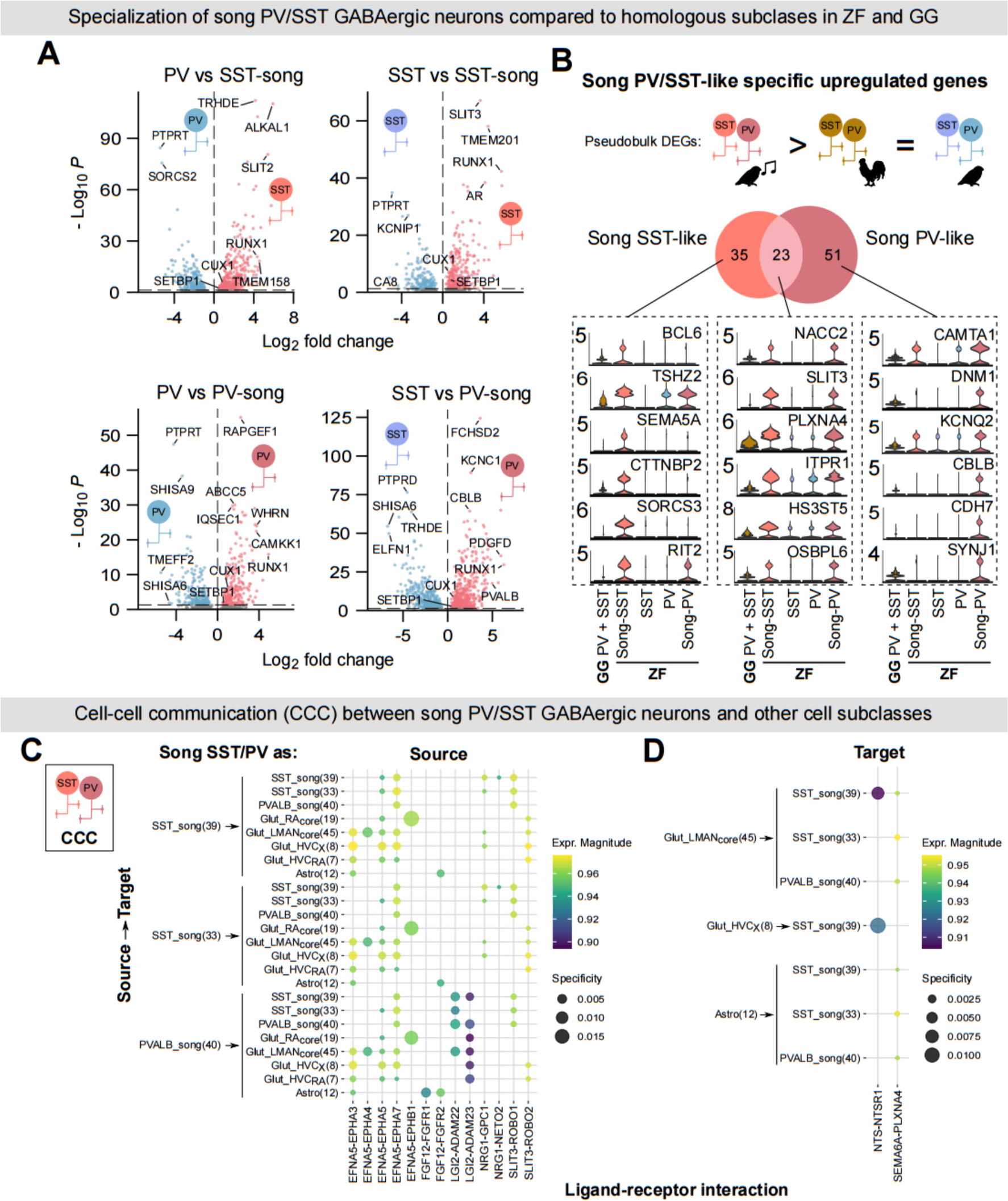
Gene expression specialization of vocal imitation GABAergic neurons. (A) Volcano plots showing pseudobulk differential expression between song-enriched PV-like and SST-like GABAergic neurons versus their ubiquitous PV- and SST-like counterparts. (B) Genes specifically upregulated in song-enriched PV- and SST-like GABAergic neurons compared to chicken PV-and SST-like GABAergic neurons, and not upregulated in the broader corresponding cell types. A Venn diagram depicts the overlap across comparisons, with a violin plot below showing select genes. (C) Predicted cell–cell communication showing ligand–receptor interactions in which song-PV and song-SST GABAergic neurons are the source and other cell types specific to song regions, and astrocytes are the target. (D) Same analysis as in (C), but with song-enriched GABAergic neurons as the target population.

To then identify gene programs in song-related GABAergic neurons that might underlie novel cell type specialization associated with vocal behavior, we focused on genes upregulated relative to both widespread zebra finch and chicken GABAergic neurons, and not significantly different between the latter two. The SST-song included upregulated expression of the transcription factors *BCL6* and *THSZ2*; axon guidance gene *SEMA5A*; ASD-linked genes *CTTNB2* involved in dendrite density in mice (*75*), *RIT2*, and *SORCS3*, also implicated in Alzheimer’s disease. Both subtypes showed upregulation of the transcription factors *RUNX1* and *NACC2*, the axon guidance genes *SLIT3* and *PLNAX4*, the ASD-risk-associated genes *ITPR1* and *HS3ST5*, and the singing-induced gene *OSBPL6*. Specific only for PV-song, upregulated genes included transcription factor *CAMTA1*; ASD-linked genes *DNM1* and *KCNQ2*; and genes upregulated in GABAergic neurons vulnerable in Alzheimer’s Disease *CBLB* and *CDH7* (*76*) (**Fig. 5B** and **Supplementary Table 7**). To assess whether gene programs enriched in song-specific GABAergic neurons might translate into distinct circuit-level interactions, we analyzed predicted cell–cell communication. We overlapped their differentially expressed genes and predicted ligand–receptor interactions inferred from expression (LIANA, agg_value < 0.05). Most predicted interactions were shared between PV- and SST-song GABAergic neurons and were not specific with respect to interaction partners or directionality. In contrast, we identified a specific interaction between the ligand EFNA5 and the receptor EPHB1 in the motor cortex analog glutamatergic cell type Glut_RA_core_(19), suggesting a potential specialized interaction between song-related GABAergic neurons and projection neurons controlling song production. In addition, all song-enriched GABAergic neurons showed upregulation of the axon guidance gene *SLIT3*, with predicted interactions with *ROBO1* in other GABAergic neurons and with *ROBO2* in glutamatergic neurons, suggesting a mechanism for establishing appropriate connectivity within song regions and for distinguishing glutamatergic from GABAergic neurons. Consistent with this interpretation, song-enriched GABAergic neurons were also targets of the SEMA6A–PLXNA4 axon guidance interaction originating from the premotor glutamatergic neuron Glut_LMAN_core_(45). Since these GABAergic neurons reside within the target region of Glut_LMAN_core_(45), this interaction may contribute to the formation of circuit architectures critical for vocal imitation (**Fig. 5C–D**).

### Comparative increase in glial and astrocyte abundance

Compared to neurons, we observed that glia formed a single or few clusters of canonical subtypes, with roughly equal membership across regions. Ependymal and RGCs were the only glial subtypes in which most cells originated from HVC or NCM (**Fig. 1C**). This was expected, as these two regions border the ventricle where these subtypes reside. Additionally, glia subclasses generally showed similar LISI score distributions regardless of whether they originated from song or non-song regions. However, astrocytes from song region dissections had a lower LISI score than other dissection regions (**Fig. 2B**), suggesting that astrocytes in song regions may exhibit a greater degree of specialization than other glia. Yet, the differential gene expression fell short of meeting the criteria for discrete taxonomic separation. Visualizing differential gene expression of astrocytes across areas confirmed that the differences were minimal (**Extended Data Fig. 8A** and **Supplementary Table 8**).

Since we observed minimal transcriptomic differences in glia across regions, we next asked whether differences might arise from changes in their relative abundance, as previously reported across regions and species (*15, 70*). We found that the percentage of glia was higher in vocal imitation regions, especially compared to their neighboring regions (HVC: 29.8% vs. NCL: 19.1% and NCM: 18.9%; LMAN: 40.6% vs. AN: 18.1%; RA: 60.3% vs. AA: 40.7%) (**Fig. 6A** and **Supplementary Table 8**). Accordingly, we found the glia-to-neuron ratio was higher in vocal imitation regions when compared with their counterparts (**Fig. 6B**). Next, we asked whether this increase also occurred at the level of a specific glia subtype. Indeed, when comparing the astrocyte-to-neuron ratio, we observed an increase across all regions compared to their counterparts (**Fig. 6A-B** and **Supplementary Table 8**). We then quantitatively corroborated these findings using immunohistochemistry against neuronal (NeuN), astrocyte (Sox9), and glia (Hoechst+ NeuN-) markers (**Fig. 6D-E** and **Supplementary Table 8**).

**Fig. 6.**
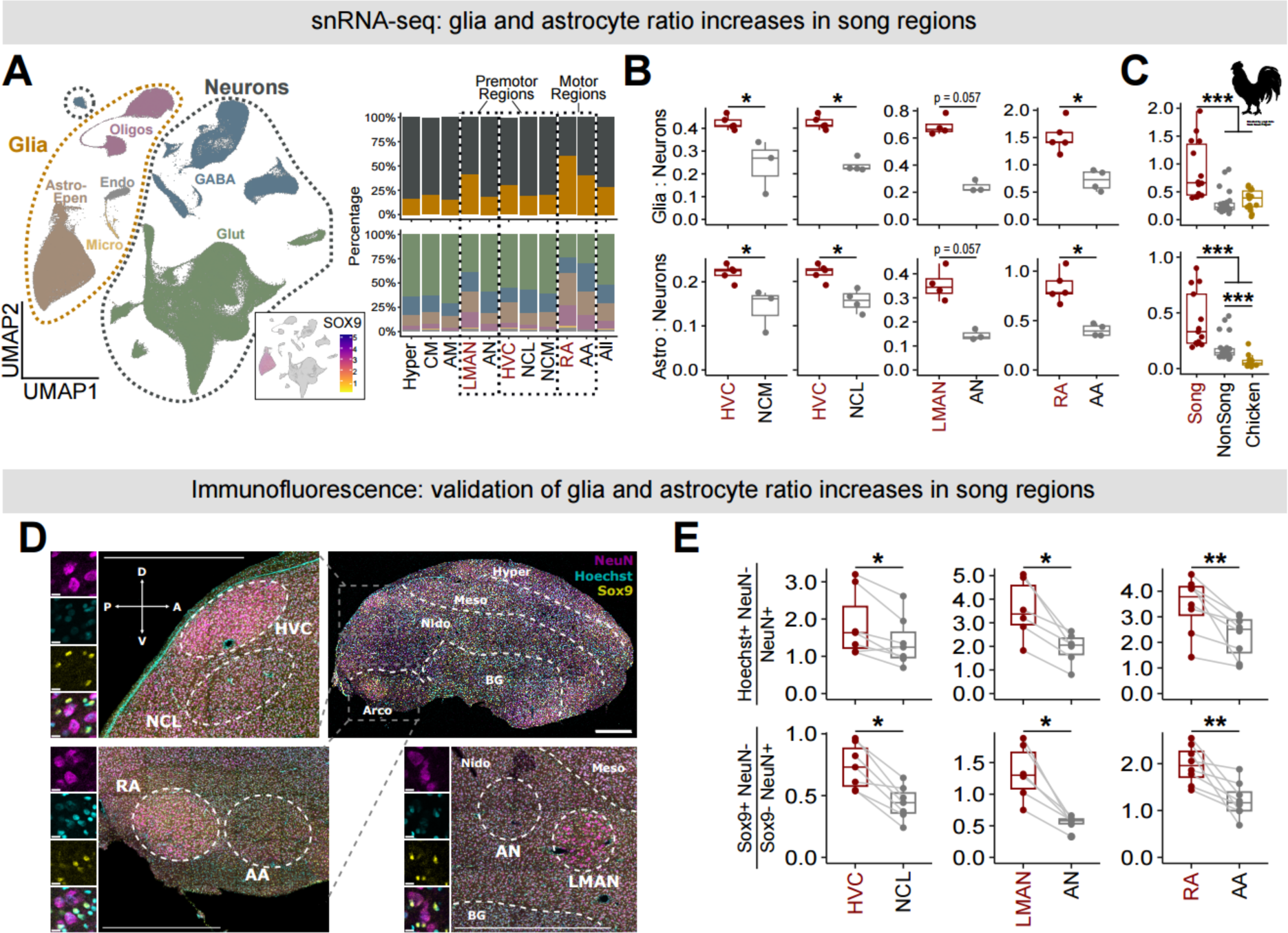
Increased astrocyte-driven glial to neuron ratio in vocal regions. (A) Left, UMAP representation of snRNA-seq colored by major cell type, with inset of *SOX9* log expression. Right, regional composition of glia/neuron (top) and major cell types (bottom). Dashed lines group song regions with anatomical counterparts. (B) The ratio from snRNA-seq of either glia to neurons (top), or astrocyte to neurons (bottom) in song regions compared to a corresponding region from the same pallial field. (C) Same ratios as in (B) grouped by song regions (red), corresponding anatomical counterparts (grey), or chicken pallium snRNA-seq dataset (yellow) (*49*). (D) Immunofluorescence (IF) analysis of neuron marker (NeuN), astrocyte (SOX9), and nuclei (Hoechst). Shown is a sagittal zebra finch brain section with enlargements of HVC, RA, and LMAN, with their corresponding control regions dissected in snRNA-seq. Each region is outlined in the enlarged images. Scale bars, 1 mm. Example cells are enlarged and split by channel. Scale bars, 10 μm. (E) Quantification of the same ratios in (B) using IF (n = 4 birds, 8 hemispheres).

Both snRNAseq and immunohistology indicated enrichment of glial cells and astrocyte support, particularly compared with neighboring areas with similar development and function. For example, although the motor regions (RA and AA) showed the highest overall glia percentages, RA had a higher ratio than AA, highlighting the importance of comparing regions with similar functions based on parallel connectivity. In this case, the comparisons indicate that the demands of vocal imitation require more glial and astrocyte support, which may not be necessary for other behaviors involving similar cell types.

We then compared these ratios to the chicken dataset. Indeed, we found that glia ratios were significantly higher in songbird vocal regions than in the homologous regions of chickens. Astrocyte ratios were likewise increased in both vocal and corresponding non-vocal imitation regions when compared to their chicken counterparts (**Fig. 6C** and **Supplementary Table 8**). Thus, pallial areas dedicated to vocal imitation exhibit elevated glial and astrocytic support relative to both non-vocal imitation regions within the same species and to species without this behavior. This suggests that the changes in pallium glial composition are an adaptation specific to vocal imitation.

## Discussion

Here, we present a transcriptomic atlas of evolutionarily novel and conserved brain regions of the bird pallium (dorsal telencephalon), spanning areas analogous to the mammalian motor, premotor, prefrontal, secondary auditory, and primary visual cortices. We focused on identifying potential cellular innovations in song-learning brain regions that underlie the eponymous vocal imitation behavior. Using within- and cross-species analyses, we identified both expected region-specific neurons and novel behaviorally specialized glutamatergic and GABAergic neurons, as well as an increased abundance of astrocytes within these vocal imitation areas.

Across vertebrates, the strongest transcriptomic signatures generally reflect developmental origin. In the case of GABAergic neurons, their domain of origin within the ganglionic eminence dictates transcriptomic similarity, and since they migrate, there is minimal difference across regions. In comparison, glutamatergic neurons remain relatively localized and thus retain strong transcriptomic signatures associated with topographic position. For example, the adjacent human primary motor and somatosensory cortex glutamatergic neurons have similar gene expression despite distinct functions (*15*). These findings so far support a general principle for pallial glutamatergic neurons: although they are more regionally distinct than other major cell classes, their gene expression is shaped more by spatial location than by behavioral specialization.

In line with this expectation, zebra finch pallial glutamatergic neurons did display the highest degree of regional specialization among major cell classes. However, neurons in the vocal-motor and vocal-premotor regions showed pronounced divergence from putative sister neuron types both within the zebra finch pallial regions in which they are embedded and in the corresponding homologous regions in chickens. A more subtle degree of specialization was observed in the song shell system, potentially reflecting its role in song learning. In contrast, glutamatergic neurons in nonvocal motor, premotor, auditory, and visual areas were comparatively conserved, showing less transcriptomic distinction from developmentally related regions.

Moreover, the mechanisms underlying this divergence appear to differ across song regions. For example, the glutamatergic neurons specific to RA (vocal motor cortex) had a greater divergence from the motor cortex of the solely innate vocalizing species and a higher proportion of non-orthologous genes compared to nidopallial song regions. This suggests that RA specialization involved not only regulatory shifts in conserved genes but also the incorporation of lineage-specific genetic elements, whereas nidopallial song regions appear to have diverged primarily through regulatory repurposing of conserved gene programs. For example, LMAN projection neurons, required for normal song development, did not clearly align with homologous cell types in chickens. Instead, it closely resembled a rare thalamic-like cluster in the chicken pallium, suggesting expansion and repurposing of a rare avian cell type for vocal learning. HVC-specific clusters diverged into distinct transcriptional identities while partially converging on mesopallial gene programs—a notable convergence given that the mesopallium predominantly forms intratelencephalic connections (*77*), is expanded in birds relative to reptiles, and contains more neurons in avian species with higher cognitive performance (*69*).

We also identified novel MGE-like GABAergic neurons, uniquely localized to vocal-imitation regions and enriched in the motor pathway. This was unexpected, given the high conservation of GABAergic cell classes across regions and species. Nonetheless, these novel SST-like and PV-like subtypes add to evidence that GABAergic divergence and innovation might bias toward MGE-derived lineages (*15, 16, 40, 49*). One of the differentially expressed genes, *ADSS1*, was also upregulated in HVC- and LMAN-projecting neurons, suggesting a shared adaptation to possibly meet high metabolic demand in vocal-imitation brain regions.

Supporting this, vocal imitation regions showed a higher glia-to-neuron ratio, primarily driven by increased astrocyte abundance. This suggests a shared requirement for enhanced glial support, which could be related to metabolic needs or higher demand for synaptic plasticity. Although astrocytes were largely homogeneous in expression across regions, our measures of regional specificity indicated that astrocytes in vocal imitation areas exhibit some degree of regional specialization.

Taken together, our findings suggest that the evolution of vocal imitation in songbirds was accompanied by the emergence of novel glutamatergic and GABAergic neuron types specifically within higher-order motor, premotor, and prefrontal cortex–analog regions. In contrast, nonvocal motor, auditory, and visual regions remained largely conserved, underscoring that cellular innovation in the songbird pallium is closely tied to the emergence of circuits supporting learned vocal behavior. The concurrent increase in glial and astrocytic abundance within these same regions further suggests that changes in both neuronal and non-neuronal cell classes may have contributed to this behavioral specialization.

Overall, these results suggest that avian pallial evolution reflects a shared tendency with mammals to reuse conserved neural architectures for diverse behaviors. However, it also demonstrates a rare case of behavior-specific cellular innovation in brain regions involved in complex behaviors. Song region specializations might be most closely compared to specializations in the human visual cortex, which has a higher degree of molecular distinction than expected based on topography alone (*15*). However, in the song system, specializations were evident even at a coarse clustering level. Across vertebrates, complex behaviors have been correlated with multiple large-scale developmental and anatomical trends—including increased neurons in the pallium and its associative cortices (*69, 78–83*), possibly from prolonged neurodevelopment (*80, 82–84*). The distinct modes of cellular innovation might represent additional mechanisms supporting behavioral innovation that is less constrained by requirements for further brain expansion. It also raises the possibility that behavior-linked cellular specialization in the pallium may be more widespread than currently appreciated, but less readily detectable in the human brain due to constraints on accessibility.

## Acknowledgments

We thank Drs. Allan-Hermann Pool, Sebastian Choi, Stephanie White, Aparna Bhaduri and members of the Roberts and Konopka laboratories for comments on initial versions of this manuscript, Jennifer Holdway, Luis Garcia, and Rico Cabuco for laboratory support, Dr. Lei Xiao for exploratory clustering of spatial transcriptomic data, and Dr. Bradley Colquitt for initial help with certain analyses. We thank Jesse Hilton, Richonda Hunte, and Pamela Jennings for administrative support.

## Funding

National Institutes of Health grant R01NS108424 (TFR) National Institutes of Health grant R01DC020333 (TFR) National Institutes of Health grant R01NS102488 (TFR, GK) National Institutes of Health grant UF1NS115821 (TFR, GK) National Institutes of Health grant F31NS131049 (CGO)

## Author contributions

Conceptualization: CGO, TFR, GK

Methodology: CGO, ZZ, AK, DPM, JJ

Investigation: CGO, ZZ, AK, DPM, JJ

Visualization: CGO

Funding acquisition: TFR, GK, CGO

Project administration: TFR, GK

Supervision: TFR, GK, CGO

Writing – original draft: CGO

Writing – review & editing: TFR, GK, CGO

## Competing interests

Authors declare that they have no competing interests.

## Data and materials availability

Data are available in the Gene Expression Omnibus for snRNA-seq (GSE316328) and the spatial dataset (GSE316807). Processed data used for analyses will be made available at https://cloud.biohpc.swmed.edu/ _____. Code used in key analyses and figure production is available at https://github.com/cgorozco/Songbird_Pallium_snRNA_sptRNA

## Declaration of generative AI and AI-assisted technologies

During the preparation of this work, the authors used OpenAI ChatGPT to support certain coding tasks and to provide feedback on drafts of some sections. The authors reviewed and iteratively edited the content as needed and take full responsibility for the final content of the publication.

## Materials and Methods

### Animal use

All zebra finches were from our breeding colonies at the University of Texas Southwestern Medical Center (UTSW) or were purchased from approved vendors. Experiments were conducted in accordance with National Institutes of Health (NIH) and UTSW policies governing animal use and welfare.

### Single-nucleus sample preparation and sequencing

Six adult male zebra finches (133-185 days post-hatch) were rapidly decapitated between 11:00 - 14:00 (4-7 hours after lights-on). Brains were dissected out and placed in ice-cold ACSF (126 mM NaCl; 3 mM KCl; 1.25 mM NaH2PO4; 26 mM NaHCO3; 10 mM D-(+)- glucose, 2 mM MgSO4; 2 mM CaCl2) bubbled with carbogen gas (95% O2, 5% CO2) and then sliced in sagittal orientation at 200 μm on a vibratome. HVC, RA, and LMAN were identified by their anatomical position and their dense myelinations. Adjacent regions of comparable size (NCL, AA, and AN) were then dissected, as were Hyper, CM, and AM, which were identified by their anatomical landmarks. Dissections were performed manually using scissors under a dissecting microscope, and tissue was transferred into tubes containing ice-cold ACSF. For each bird and region, two to three samples were collected from each hemisphere. When all tissue was collected, excess ACSF was removed from each tube, samples were flash-frozen in liquid nitrogen, and stored at -80 °C.

For nuclei extraction, the tissue was thawed over ice and then homogenized in 500 µL of ice-cold Nuclei EZ lysis buffer (EZ PREP NUC-101, Sigma) using a pre-chilled glass Dounce homogenizer. Debris was removed via density gradient centrifugation using Nuclei PURE 2 M sucrose cushion solution and Nuclei PURE sucrose cushion buffer (Nuclei PURE prep isolation kit, no. NUC201-1KT, Sigma Aldrich). These two solutions were mixed in a 9:1 ratio, and 500 µL of the sucrose solution was added to a 2-ml Eppendorf tube. The homogenized nuclei suspension was mixed with 900 µL of sucrose cushion solution by pipetting 10 times, resulting in a 1,400 µL volume, which was carefully layered over the sucrose buffer without mixing. The samples were centrifuged at 13,000 x g for 45 min at 4 °C, and all but 100 µL of supernatant was discarded. Nuclei were resuspended in 300 µL of nucleus suspension buffer (NSB), consisting of 1× PBS, 2% BSA (no. AM2618, Thermo Fisher Scientific), and 0.4 U µL −1 RNAse inhibitor (no. AM2694, Thermo Fisher Scientific). Samples were then centrifuged at 550 x g for 5 min at 4 °C, and all but 50 µL of supernatant was discarded. The nuclear pellet was resuspended in NSB and filtered through a 40-μm Flowmi cell strainer (Bel-Art Products, #H13680-0040) into a new tube. The nuclei concentration was measured using Hoechst and decreased to 1,000 nuclei per µL with NSB if necessary.

Droplet-based snRNA-seq libraries were prepared using the Chromium Single Cell 3′ v3.1 (10x Genomics) kit, following the manufacturer’s protocol. Amplified cDNA and final libraries were quality-controlled using a TapeStation and quantified by Qubit. Libraries were sequenced on an Illumina NovaSeq 6000. Samples that did not pass a threshold of 1,000 median genes detected per cell were re-sequenced to increase sequencing depth. Additionally, raw snRNA-seq data from zebra finch samples published by Colquitt et al. (*40*) were reprocessed like the data here and combined with data from our own study. Except for one of the HVC libraries, which failed to integrate with the rest of the dataset.

### Single-nucleus preprocessing

Raw sequencing data were obtained as binary base calls (BCL files) from the sequencing core. The *mkfastq* command from 10X Cellranger v.7.2.0 or *bcl2fastq* command from Illumina v2.20.0 was used to demultiplex the BCL files. Paired-end FASTQ files (28 bp long R1—cell barcode and UMI sequence; 90 bp long R2—transcript sequence) were aligned to a reference zebra finch genome (bTaeGut1.4.pri). Then the *count* command from 10X Cellranger v.7.1.0 was used to count reads as the number of unique molecular identifiers (UMIs) per gene per cell. The reference genome and annotation index for pre-mRNAs were used to account for reads aligned to both introns and exons.

To remove ambient RNA contamination, we used CellBender on the unfiltered output from Cellranger per sample (*85*). The empty-droplet-filtered output from CellBender was used in further processing. Each sample was then clustered with the Seurat standard pipeline and clustered [SCTransform(), RunPCA(npcs=50), FindNeighbors(dims= 1:30), RunUMAP(dims =1:30), FindClusters(resolution=2)], followed by removal of clusters with low intronic read ratios using the DropletQC package(*86*), i.e., the average and median of the nuclear fraction reads were lower than 0.5. Additionally, each library was evaluated individually using standard Seurat pipelines to assess overall data quality, and libraries that failed to produce well-defined clusters were excluded from further analysis. From the remaining libraries, we kept only nuclei with >200 UMIs and mitochondrial read mapping <5%.

Each library was then processed with scDblFinder(*87*) to identify and score putative doublets. Nuclei classified as singlets were retained, and doublet scores were incorporated with additional annotation-based criteria to identify residual doublets. Broad cell-class annotations were obtained using Azimuth(*88*) (using the mouse motor cortex reference(*89*)) and CZ CELLxGENE(*90*) (pipeline was developed by UTSW BioHPC), and were used to aid doublet classification. Likely doublets were identified by co-expression of marker genes typically expressed in different cell classes. These were removed through iterative standard clustering and filtering that incorporated all the described criteria until no such clusters remained.

### snRNA-seq data integration and clustering

The nuclei were then re-clustered following the quality-control iterations (see above). Each library was normalized and scaled using the SCTransform function. Nuclei with outlying numbers of detected genes were identified within neurons and glia separately using an interquartile range–based threshold and removed. Samples were then integrated using reciprocal principal component analysis (RPCA), and a k-nearest neighbors graph was constructed in the first 30 principal components of this integrated space.

Using the Allen Institute’s scrattch.hicat package, we performed 50 clustering iterations in principal component space with 80% of random subsampled cells to produce a consensus of high-confidence clusters (*43, 44*). We used the recommended default parameters: padj.th = 0.05, lfc.th = 1, low.th = 1, q1.th = 0.3, q.diff.th = 0.7, and de.score.th = 150. The normalized data was then averaged within each cluster, and the differentially expressed genes identified during clustering were used for building the hierarchical dendrogram. To annotate cell clusters, we relied on expression of canonical marker genes (such as GAD1 for GABAergic neurons), hierarchical clustering, predicted annotations, and the proportion of nuclei belonging to each anatomical dissection.

### Technical note on identifying the ‘shell’ regions surrounding vocal imitation areas

We distinguished between song-region-specific and surrounding (shell’) glutamatergic subtypes using a variety of complementary methods. This included spatial mapping of the snRNA-seq clusters to the sptRNA-seq (see Visium analysis); fluorescence in situ hybridization targeting cluster-specific marker genes, along with retrograde tracers to guide anatomical mapping; and expression of marker genes identified by previous findings (*26, 38, 41*).

In situ expression of the Glut_LMANcore(45) upregulated genes *CALCB* and *ADSS1* were confined to LMAN projection neurons (retrograde tracer), exhibiting complete overlap with SLC17A6⁺ (glutamatergic neuron marker). In contrast, Glut_LMANshell(44), which also had the majority cells originating from the LMAN dissection, lacked *CALCB* and *ADSS1* expression (**Fig. 2D** and **Extended Data Fig. 3A-B**). One possibility was that these cells originated from the LMAN shell, an area that forms part of the broader “shell system” that immediately surrounds and thus closely parallels the vocal imitation circuitry (*27–35*). Supporting this interpretation, Glut_LMANshell(44) expresses *SEMA5A*, an axon guidance gene upregulated in both LMAN core and shell, but not *CNN3*, which is only enriched in the core. It was also positive for genes downregulated in the core, *CAMK2A* and *UNC5D*. Accordingly, the spatial mapping showed Glut_LMANcore(45) and Glut_LMANshell(44) positioned in proximity but not completely overlapping (**Extended Data Fig. 3A-B**).

Similarly, clusters Glut_RAcup(18) and Glut_ArcoPall(27), which had many cells coming from the RA dissection, were negative for *SRD5A2* and *SLC4A11*, genes with restricted expression to RA(*41*), and instead expressed downregulated genes in RA, *TENM3* and *CNTN5* (**Extended Data Fig. 3C-D**). Likewise, Glut_HVCshelf(28), from which many cells originated from HVC, lacked expression of *ADSS1*, a gene expressed in both Glut_HVC-RA(7), Glut_HVC-X(8), and validated with in situ to be highly restricted to and co-labeling all SLC17A6⁺ neurons in HVC. Cluster Glut_HVCshelf(28) also did not express the HVC upregulated genes *SEMA3E* and *WNT5B*. Although it expressed *ALDH1A2* and *CADPS2*, genes upregulated in both HVC and HVC shelf, it also expressed the HVC downregulated genes *GRIA4* and *CACNA1G* (**Extended Data Fig. 3E-G**).

Despite forming distinct clusters, these shell pathway clusters contained cells from non-vocal-imitation region dissections. This suggests that the shell system may have less discrete boundaries, where neighboring dissected regions converge into a continuum with them. Alternatively, the shell system might share greater transcriptional similarity with surrounding regions or simply encompass a larger population of related glutamatergic neurons. Nonetheless, by leveraging these spatially restricted marker genes, we were able to distinguish the projection neurons within the vocal imitation regions from the closely associated glutamatergic populations of the surrounding shell system.

The gene expression profile of Glut_RAcore(19) identified it as the sole glutamatergic cluster corresponding to neurons of the vocal motor cortex. Likewise, Glut_LMANcore(45) represents the core population of projection neurons within the vocal premotor cortex, LMAN. In addition, Glut_HVC_RA_(7) and Glut_HVC_X_(8) corresponded to two distinct, non-overlapping projection neuron types that target the vocal motor cortex RA and vocal basal ganglia (Area X), respectively, and which are distinguished by the known marker genes *NTS* and *UTS2B*.

### Differential gene expression

For single-cell-based differential gene expression (DGE) analysis, we used the Wilcoxon rank-sum test to visualize DGE in GABAergic neurons and astrocytes. Differentially expressed genes were defined as those with an adjusted p-value < 0.05, a log fold-change exceeding ±0.1, and a minimum percent difference of 15.

All pseudobulk differential gene expression was carried out using DESeq2 (*91*) and were considered significant if the adjusted p-value was < 0.05.

For GABAergic neurons, nuclei from which fewer than 20 nuclei from a given zebra finch contributed to a cluster were excluded(*92*). From the remaining nuclei, gene-expression counts were aggregated across all cells within each cluster and across individual birds to generate pseudobulk replicates. Differentially expressed genes were defined as those with a log fold change of at least ±0.1.

Comparisons among zebra finch GABAergic neurons were performed at the subclass level and between those enriched in vocal-imitation regions and the rest. Specifically, song-enriched GABAergic neuron subclasses (i.e., PVALB-song or SST-song) were compared against their corresponding widespread GABAergic neuron subclasses (i.e., other PVALB or SST).

For comparisons to chicken GABAergic cell types, analyses were restricted to one-to-one orthologous genes. Zebra finch song-enriched, and widespread GABAergic neuron subclasses were each compared to a reference group comprising all SST- and PV-like GABAergic neurons in chicken, with chicken nuclei aggregated within each cell cluster and individual. Chicken SST-and PV-like GABAergic neurons were grouped for comparisons. PV/SST song-specific upregulated genes were defined as those significantly upregulated in zebra finch song-region-enriched subclasses relative to both widespread zebra finch GABAergic neurons and chicken GABAergic neurons, while showing no significant differential expression between widespread zebra finch GABAergic neurons and chicken GABAergic neurons (i.e., ZF-song > ZF-other = CHK, where “=” denotes no significant differential expression).

For pseudobulk differential gene expression analysis of glutamatergic neurons, nuclei from which fewer than 2% of nuclei from a given dissection region contributed to a cluster were excluded (*92*). From the remaining nuclei, gene-expression counts were aggregated across all nuclei within each cluster and across individual birds to generate pseudobulk replicates. Differentially expressed genes were defined as those with a log fold change of ±0.1 or greater and expression in all pseudo-replicates of at least one compared cluster. Within each pallial region, each cluster was compared pairwise with all others, and the resulting differentially expressed genes were consolidated for each cluster. The differentially expressed genes were then used to generate the upset plots.

In the pallial region comparisons, in addition to the 2% threshold, song regions were excluded. The remaining nuclei expression counts were aggregated across all nuclei within each pallial region (i.e., Hyperpallium, Mesopallium, Nidopallium, Arcopallium) and across individual birds to generate pseudobulk replicates. Differentially expressed genes were defined as those with a log fold change of ±0.2 or greater and expression in all pseudo-replicates of at least one compared pallial region. The differentially expressed genes were then used to generate the upset plots. Genes upregulated in a given pallial region relative to at least one other region were defined as that region’s module genes and used to score each cluster.

### Local inverse Simpson’s index analysis

The degree of transcriptomic cell type similarity across dissection regions was quantified using the local inverse Simpson’s index (LISI) (*48*). This metric is independent of the resolution of the integrated clusters. It identifies Gaussian-kernel–based local neighborhoods for each cell in a shared, integrated feature space and estimates the average number of draws needed to sample another cell with the same batch label from that neighborhood. A LISI value of one indicates no mixing between datasets, whereas higher values reflect higher mixing within the embedded transcriptomic space. Each cell has a LISI score, which can be grouped into different categories to analyze mixing at multiple resolutions.

LISI scores were computed in the PCA space derived from the integrated assay using the first 30 principal components, providing a measure of mixing across the ten dissected regions. Scores were grouped and compared between vocal imitation regions and other regions at the cell class level. For cell–type–level analyses, cell types contributing fewer than 2% of glutamatergic neurons and 5% in in GABAergic neurons within a given cluster were excluded. Scores were then averaged by bird and grouped by cluster. Lower LISI scores were interpreted as greater regional transcriptomic specificity, whereas higher scores indicated greater mixing between regions in latent space and thus increased transcriptomic similarity.

### Cell-type fraction and ratio comparisons

For comparison of cell-type ratios, we calculated the fraction of GABAergic subtypes within all GABAergic cells, the ratio of glial to neurons, and the ratio of astrocytes to neurons for each individual bird in snRNA-seq and for each brain hemisphere in situ and IF. To determine whether fraction or ratio differences were significant between comparable regions, we calculated the P value using Wilcoxon Rank-Sum test.

### Cell–cell communication analysis

Cell–cell communication events were predicted using the LIgandreceptor ANalysis frAmework (LIANA) in R(*93*). Specifically, the ligand–receptor analysis was performed using the liana_wrap() function with the default methods, which incorporates the resources NATMI(*94*), Connectome (*95*), SingleCellSignalR (*96*), and CellPhoneDB (*97*). We then ran liana_aggregate() to find consensus ranks between the different methods. Only interactions (ligand–receptor pairs) with a robust rank aggregation (RRA) (*98*) score smaller than 0.05 (aggregate_rank < 0.05) were considered in downstream analyses. To highlight possible upregulated ligand–receptor interactions in cell types of interest, we overlapped these significant interactions with differentially expressed genes of relevant comparisons.

### Gene specificity scoring

Genes whose expression had high cluster specificity were identified using the approach described in (*40, 49, 51*). For each gene, a specificity score was calculated as ∑ *log* (*x*_n_ / *x̅*_n_), where *x*_n_ is each gene’s expression divided by the sum of expression across all clusters and x̅_n_ is the mean of this value.

### Cross-species correlation

We compared cell type populations across species using two complementary approaches: Gene Specificity Index (GSI) correlations (*51*) and Seurat’s canonical correlation analysis (CCA)–based integration (*50*), both restricted to sets of one-to-one orthologous genes, similar to our previous approach (*40*). To assign an overall similarity metric, scores from each method were summed, resulting in a possible composite similarity score ranging from 0 to 2, similar to Zaremba et al., 2025. Each method included its own permutation-based significance testing, such that the aggregated similarity score could reflect zero, one, or two significant correlations.

For the GSI-based analysis, differentially expressed genes for each dataset in the comparison were identified using Seurat FindAllMarkers (Wilcoxon Rank Sum, min.pct = 0.2 logfc.threshold = 0.1, min.diff.pct = 0.3). For each differential expression list, the 50 significant genes (adjusted p-value < 0.05) with the highest fold-change in each cluster were selected. Average cluster expression profiles were then subset to this shared gene set and transformed by log(x+1) + y, where y was a small scalar (0.1). A specificity score was calculated in a manner similar to Tosches et al., 2018 (*51*), such that expression values were divided by the across-cluster mean for each gene. The resulting specificity matrices (songbird and chicken) were then compared by computing Spearman correlations between corresponding cluster columns. Negative correlations were set to zero and excluded from significance testing. Significant correlations for GSI were identified using a permutation test, in which gene labels were shuffled for one of the matrices, then the two matrices were correlated using Spearman correlations (500 shuffles). Observed correlations greater than 99.5% of the shuffled distribution were considered significant.

For label transfer, comparisons were restricted to variable genes shared between cell classes being compared in each dataset (e.g., zebra finch and chicken), identified using SCTransform (variable.features.n = 6000). Cluster similarity was computed by first identifying cross-dataset anchors using FindTransferAnchors (reduction = “cca”), followed by label prediction using TransferData with default parameters. Cluster-level similarity was then quantified as the mean prediction probability for each cluster pair. Significance was similarly assessed with permutation testing, in which cell-type labels in the reference dataset were shuffled before label transfer (300 shuffles). Observed similarities exceeding 99.95% of the shuffled distribution were considered significant.

### RNA fluorescence in situ hybridization

Birds were perfused with 4% paraformaldehyde (PFA) in phosphate-buffered saline (PBS). Brains were postfixed overnight in 4% PFA at 4 °C with gentle agitation, then cryoprotected by incubation in 30%(g/g) sucrose prepared in the same fixative at 4 °C overnight. Brains were sectioned at 30 µm thickness and sections were collected into 0.5–1% PFA (4% PFA diluted in RNase-free PBS).

Hybridization chain reaction (HCR) RNA fluorescence in situ hybridization (FISH) was performed using a protocol modified from Ben-Tov et al., 2023(*99*). Free-floating sections were washed twice in PBS (3 min each) and incubated in 5% SDS in PBS for 45 min. Sections were rinsed twice in 2× saline-sodium citrate with 0.1% Tween-20 (2× SSCT) and incubated in 2× SSCT for 15 min with gentle agitation. Sections were pre-incubated in 300ul probe hybridization buffer for 5 min, then transferred to probe solution (1.2% final concentration) containing target-specific probe sets and incubated for 24 h at 37 °C in the dark. For multiplex experiments, probe sets carrying distinct HCR initiator tags were combined in the same hybridization reaction.

Following hybridization, sections were washed four times for 15 min each in pre-warmed probe wash buffer at 37 °C, followed by two washes in 2× SSCT (5 min each) at room temperature. Sections were then incubated in HCR amplification buffer for 30 min at room temperature. Fluorescent hairpins corresponding to each initiator tag were heat-denatured at 95°C for 90 s, cooled to 4 °C, and equilibrated in the dark for 30 min prior to use. Sections were incubated in amplification buffer containing fluorescent hairpins (1.5% final concentration) for 24 h at room temperature with gentle agitation.

After amplification, sections were washed four times in 2× SSCT for 15 min each at room temperature, mounted onto slides, air-dried briefly, and coverslipped with Fluoromount-G for imaging. Composite images were acquired and stitched using an LSM 880 laser scanning confocal microscope (Carl Zeiss, Germany).

### FISH quantification

Image quantification for various probes was performed manually using Fiji/ImageJ(*100*). Regions of interest were identified anatomically, and retrograde tracers (Fast Blue) further helped distinguish vocal imitation regions. Tracers were injected into RA for LMAN and HVC, or into hindbrain vocal circuits for RA. For cell counting, 2-3 sub-areas (50 μm x 50 μm) were randomly selected within each region of interest to quantify cell numbers. Results were averaged between these subareas and then across 2-3 slices per brain hemisphere. Ratios were quantified based on the gene expression marker of interest. For example, GAD1+ expression was used to identify GABAergic neurons, and retrograde labeling and anatomical criteria were used to determine whether they were within the vocal imitation region. The fraction of those cells that overlapped with the expression of different markers, such as *ADSS1*, was then counted.

### Immunofluorescence (IF)

Birds were perfused with 4% paraformaldehyde (PFA) in phosphate-buffered saline (PBS). Brains were postfixed overnight in 4% PFA at 4 °C with gentle agitation, then cryoprotected by incubation in 30%(g/g) sucrose prepared in the same fixative at 4 °C overnight. Brains were sectioned at 30 µm, and the sections were collected in PBS.

Fixed sections were washed 3 times for 10 minutes in PBS and then blocked in 5% normal donkey serum in PBST (0.3% Triton X-100 in PBS) for 1 hour at RT. After one 10-minute wash in PBS, slices were incubated with primary antibodies diluted in the blocking buffer (5% donkey serum in PBST) at 4°C for 24 hours.

Slices were then washed 3 times for 10 minutes in PBS and incubated with fluorescent secondary antibodies and Hoechst (diluted in blocking buffer) at RT for 1 hour. After a final PBS wash, sections were mounted onto slides with Fluoromount-G (eBioscience, CA, USA). Composite images were acquired and stitched using an LSM 880 or LSM 710 laser scanning confocal microscope (Carl Zeiss, Germany). The following primary antibodies were used: mouse anti-NeuN (Sigma-Aldrich MAB377; 1:1000) and rabbit anti-Sox9 (Millipore AB5535; 1:1000). The following secondary antibodies were used: donkey anti-mouse conjugated to Alexa Fluor 568 (Invitrogen A10037; 1:1000) and donkey anti-rabbit conjugated to Alexa Fluor 488 (Invitrogen A21206; 1:1000).

All image analyses were performed using Fiji/ImageJ(*100*) and quantification was done the same as for FISH.

### Visium tissue processing and sequencing

Spatial transcriptomic profiling was performed using the 10x Genomics Visium Spatial Gene Expression platform, following the manufacturer’s protocol. Briefly, fresh tissue was rapidly harvested and immediately flash frozen by immersion in pre-chilled isopentane cooled with liquid nitrogen. Embedded tissue blocks were stored at −80 °C until further processing.

Cryosectioning was performed on a cryostat maintained at −20 °C. Tissue blocks were equilibrated in the cryostat prior to sectioning, and sections were cut at a thickness of 10 µm. Regions containing HVC, RA, and LMAN were identified by their anatomical position and dense myelination. Sections were carefully mounted onto Visium Spatial Gene Expression Slides (10x Genomics), ensuring placement within the capture area, and immediately returned to −80 °C for short-term storage or processed directly. A total of 24 sections from three adult male zebra finches (140-145dph) were collected, seven of which were used in this study based on quality control and the presence of regions of interest.

Prior to library preparation, tissue sections were fixed in chilled methanol and stained with hematoxylin and eosin (H&E) following the Visium protocol. Permeabilization time was optimized according to tissue type using the Visium Tissue Optimization workflow (18 min), after which reverse transcription was carried out in situ to generate spatially barcoded cDNA.

Amplification of cDNA and library construction were performed according to the 10x Genomics Visium Spatial Gene Expression User Guide. Amplified cDNA and final libraries were quality-controlled using a TapeStation and quantified by Qubit. Libraries were sequenced on a NovaSeq 6000 and one section on a NextSeq500, using paired-end sequencing with read lengths consistent with Visium specifications (Read 1: 28 bp, i7 index: 10 bp, i5 index: 10 bp, Read 2: 90 bp).

Raw sequencing data were processed using Space Ranger v1.2.2 with default parameters. Reads were aligned to the reference zebra finch genome bTaeGut1.4.pri, and spatial gene expression matrices were generated for downstream analyses.

### Visium analysis

Individual samples were first quality-controlled with a standard Seurat clustering workflow, with slices of poor quality or lacking regions of interest excluded. Each remaining library was then normalized using SCTransform, with the number of genes and transcripts regressed during normalization, and highly variable genes identified. Samples were integrated using reciprocal principal component analysis (RPCA) (*50*) and clustered at low resolution to distinguish the pallium from other brain regions. The normalized data were used for visualizing transcript expression.

We then used canonical correlation analysis (CCA)–based integration (*50*) to predict the locations of our pallium snRNA-seq clusters in the spatial transcriptomics data. Furthermore, the low-resolution clustering, combined with minimal manual selection guided by anatomy, allowed the identification of pallium-specific beads. For clarity and relevance, only the pallium-specific beads were used for visualization of spatial prediction scores.

---

**Extended Data Fig. 1.**
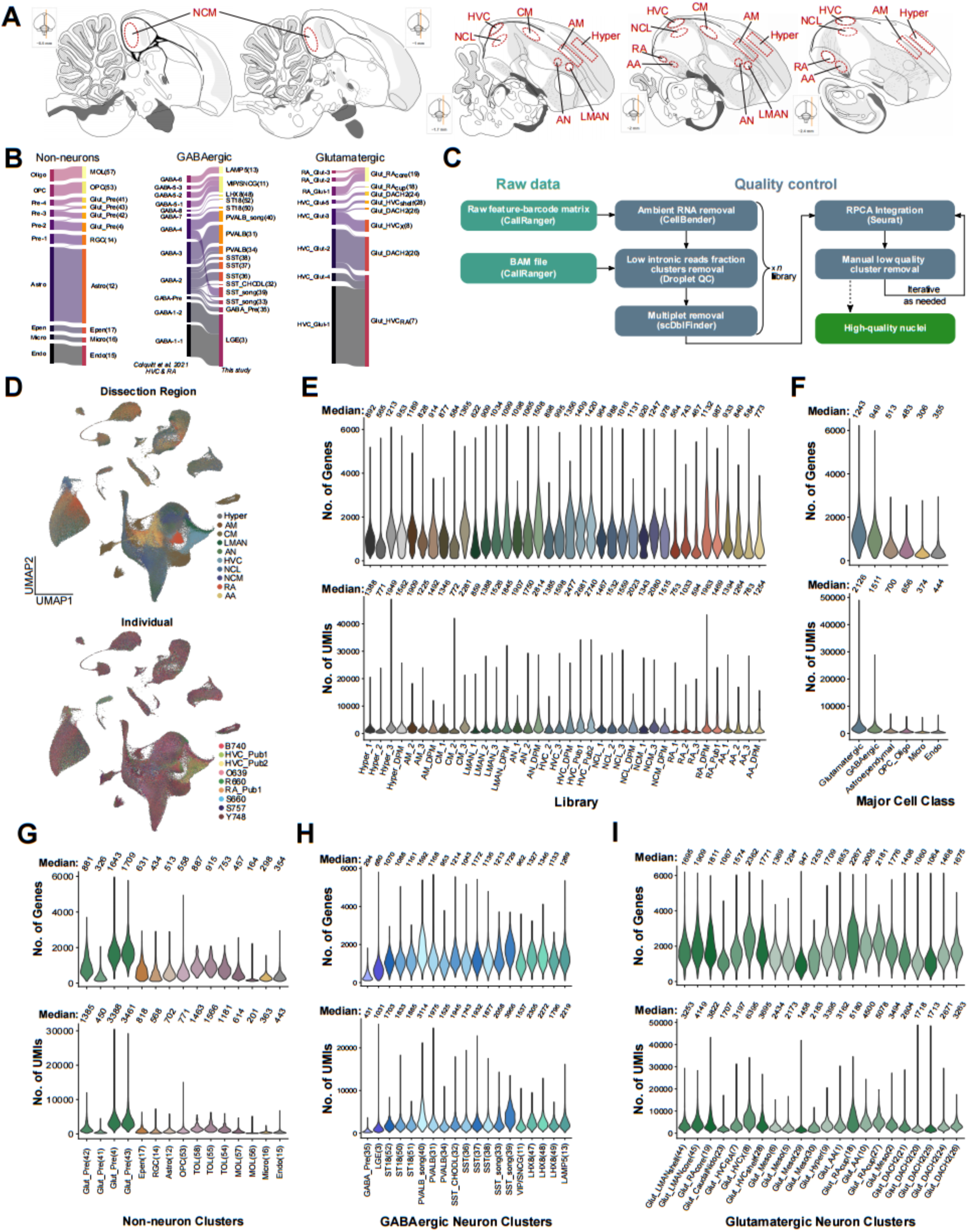
snRNAseq dataset collection and quality control. (A) Targeted zebra finch dissection regions (red outlines) across sagittal sections based on the ZEBrA database brain atlas (Oregon Health & Science University, Portland, OR 97239; http://www.zebrafinchatlas.org). (B) New cluster annotation of previously published HVC and RA snRNA-seq dataset (zebra finch only) (*40*) to this study. (C) Quality control pipeline for the snRNA-seq dataset to ensure high-quality nuclei. (D) UMAP of zebra finch pallium dataset colored by dissection region, top, or sampled individuals, bottom. (E) Number of detected genes, top, and UMI counts, bottom, per snRNA-seq library after selection of high-quality nuclei, gradient colored by dissection region. (**F – I**) Same as (E), per grouping of major cell class (F), non-neuron clusters (G), GABAergic neuron clusters (H), or glutamatergic neuron clusters (I), colored by cell class/cluster.

**Extended Data Fig. 2.**
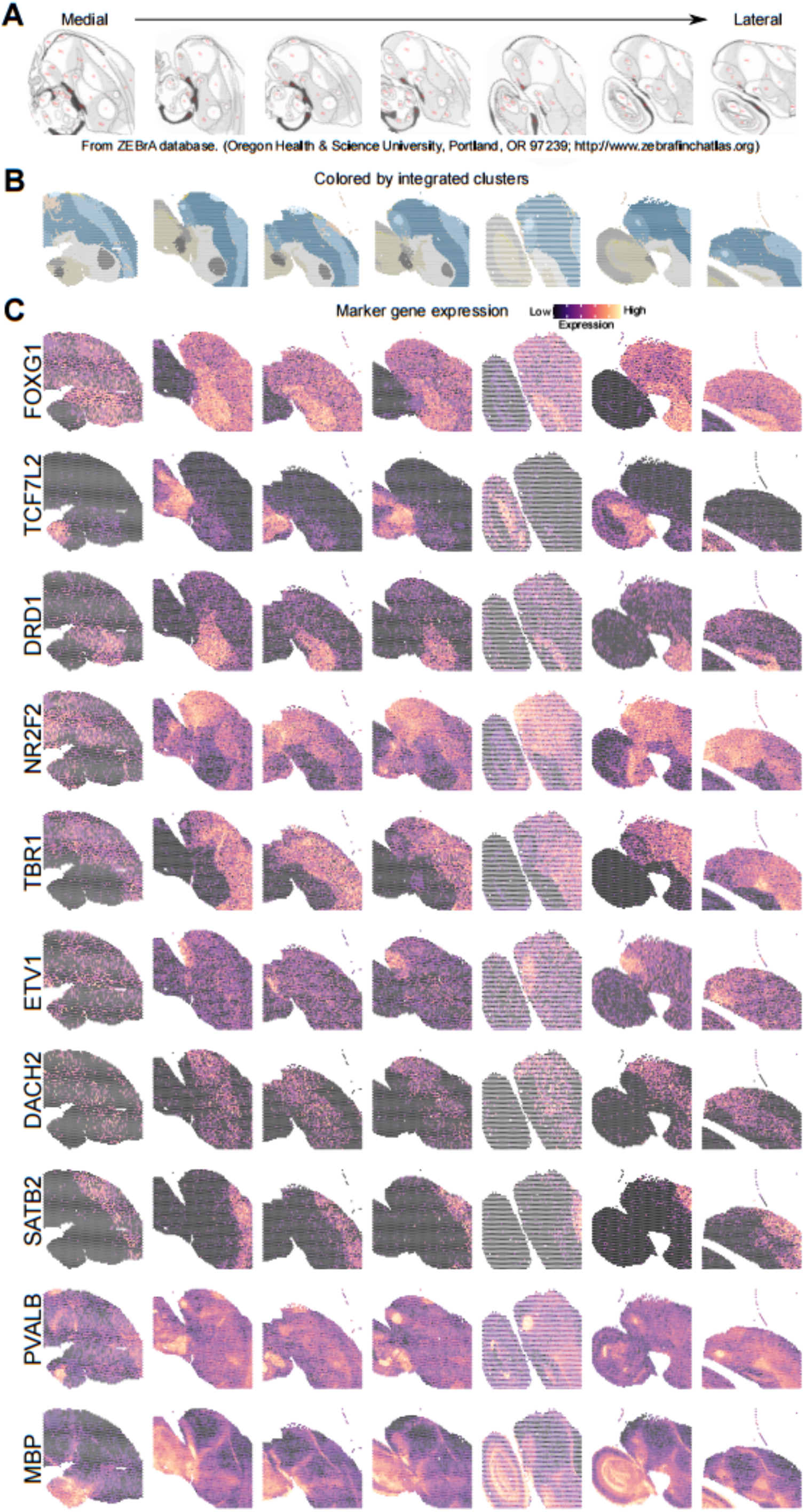
Spatial transcriptomic clusters and gene expression consistently reflect anatomical organization. (A) Zebra finch brain atlas aligned approximately to the corresponding sptRNA-seq brain slices, ordered roughly from medial to lateral. Acronyms for annotated anatomical regions are as in zebrafinchatlas.org. (B) Integrated clusters and (**C**) expression of regional marker genes (as in Fig. 1F) across all sagittal sections.

**Extended Data Fig. 3.**
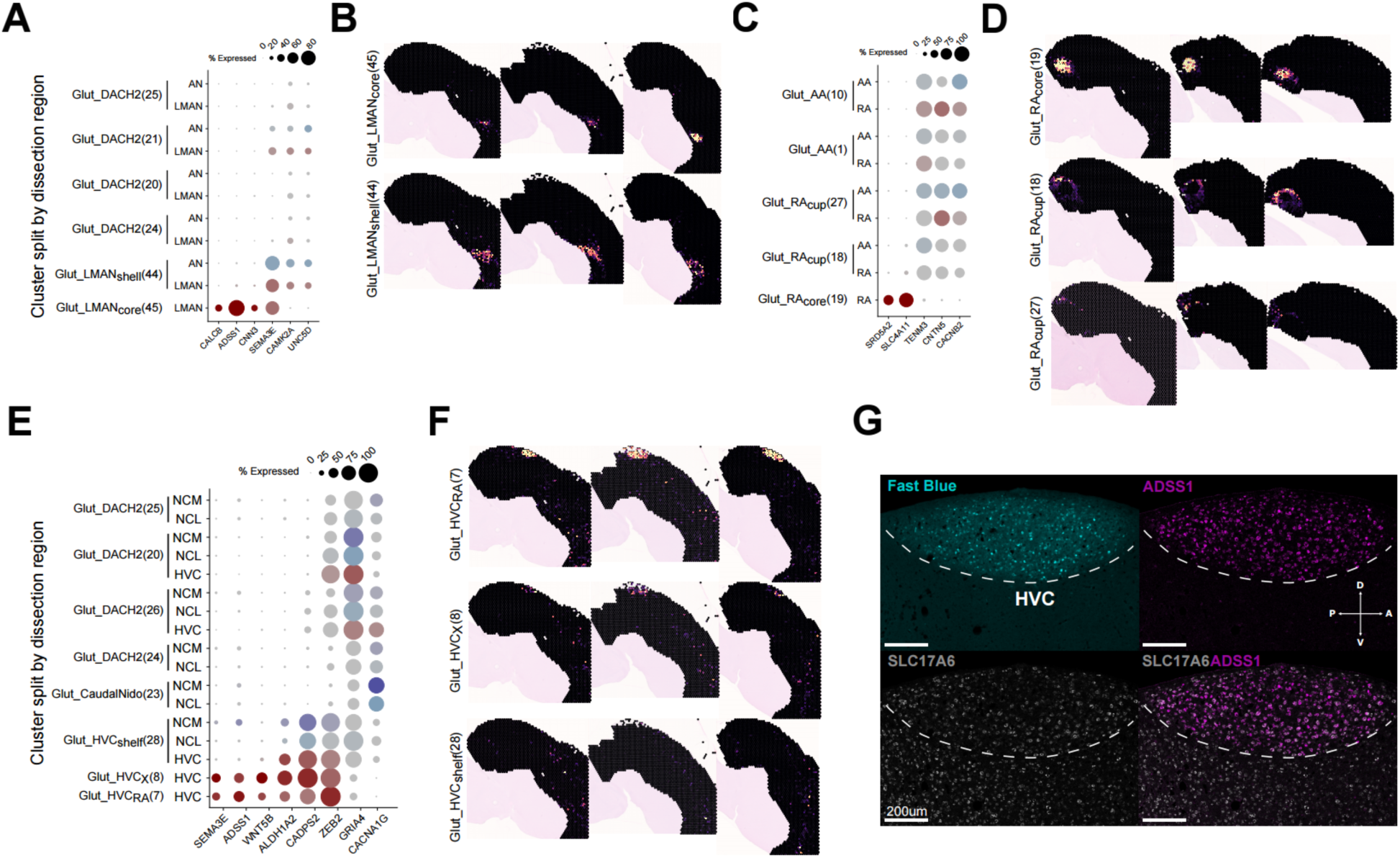
**Identification of the song shell system.** (A) Gene expression dotplot of clusters split by region, excluding cells from any region that contributed less than 2% of the total cells in that cluster. Shown are select genes upregulated in LMAN (*CALCB*, *ADSS1*, *CNN3*), both core and shelf (*SEMA3E*), or downregulated in LMAN (*CAMK2A*, *UNC5D*). Dot size indicates the percentage of cells expressing, and the darker the color, the higher the expression (red for HVC and blue for the other regions). (B) Spatial prediction of clusters that had most of their cells with membership to the LMAN dissection. (C) Same as (A), but genes upregulated in RA (*SRD5A2*, *SLC4A11*) or downregulated in RA (*TENM3*, *CNTN5*, *CACNB2*). (D) Same as (B), but with most of their cells with membership to the RA dissection. (E) Same as (A), but genes that are upregulated in HVC core (*SEMA3E, ADDS1*, *WNT5B*), both core and shelf (*ALDH1A2*, *CADPS2*, *ZEB2*), or downregulated in HVC (*GRIA4*, *CACNA1G*). (F) Same as (B), but with most of their cells with membership to the HVC dissection. (G) Split channels from a sagittal section of a bird injected with Fast Blue in RA to identify HVC; combined with *in situ* hybridization for one of HVC’s marker genes (*ADSS1*) or glutamatergic neuron marker (*SLC17A6*), and merged image.

**Extended Data Fig. 4.**
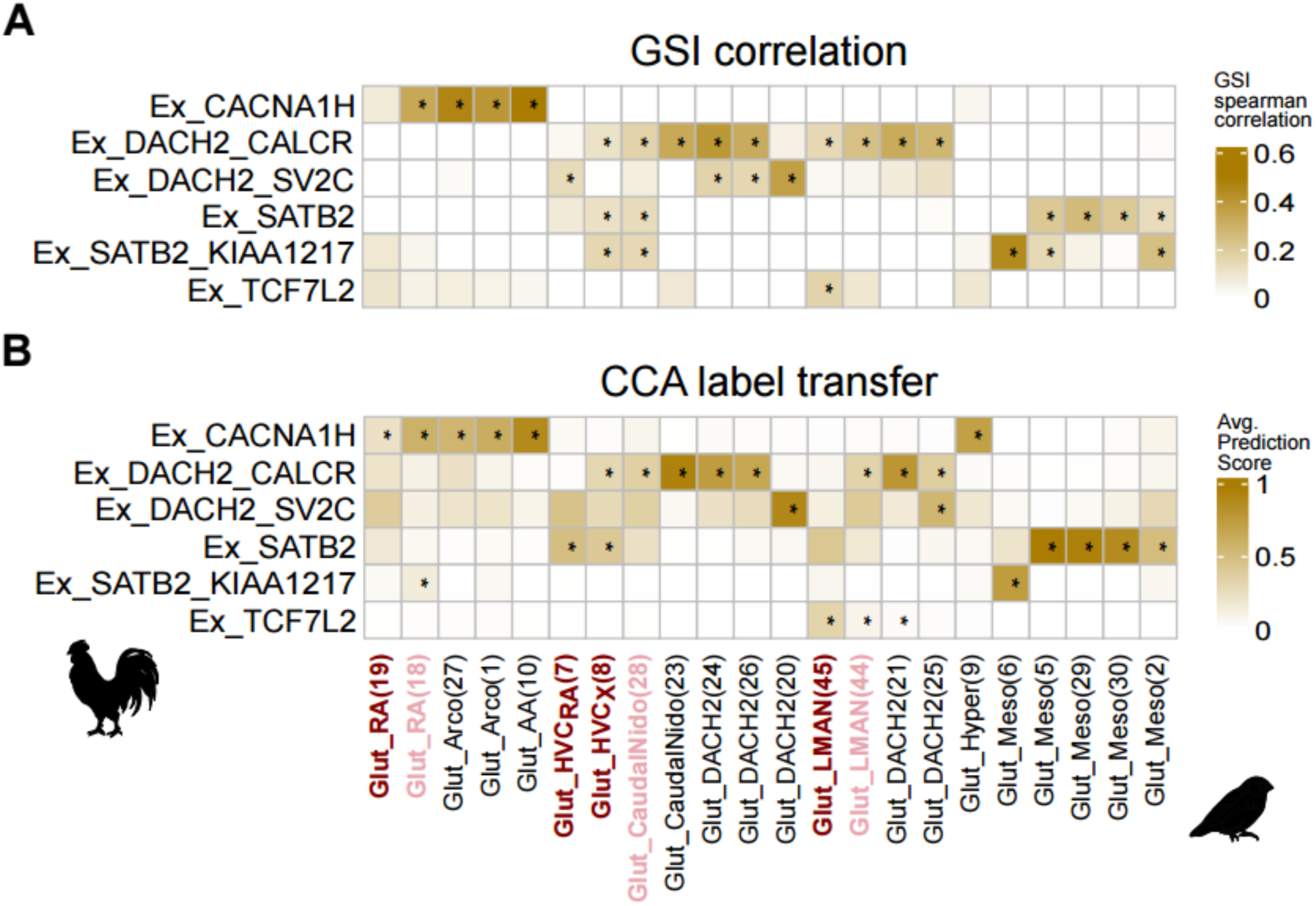
Comparison of glutamatergic neurons between chicken and zebra finch pallium. Comparison between glutamatergic supertypes in the chicken pallium to clusters of glutamatergic neurons in the zebra finch pallium based on either (**A**) gene specificity index, Spearman correlation, or (**B**) label transfer average prediction score method.

**Extended Data Fig. 5.**
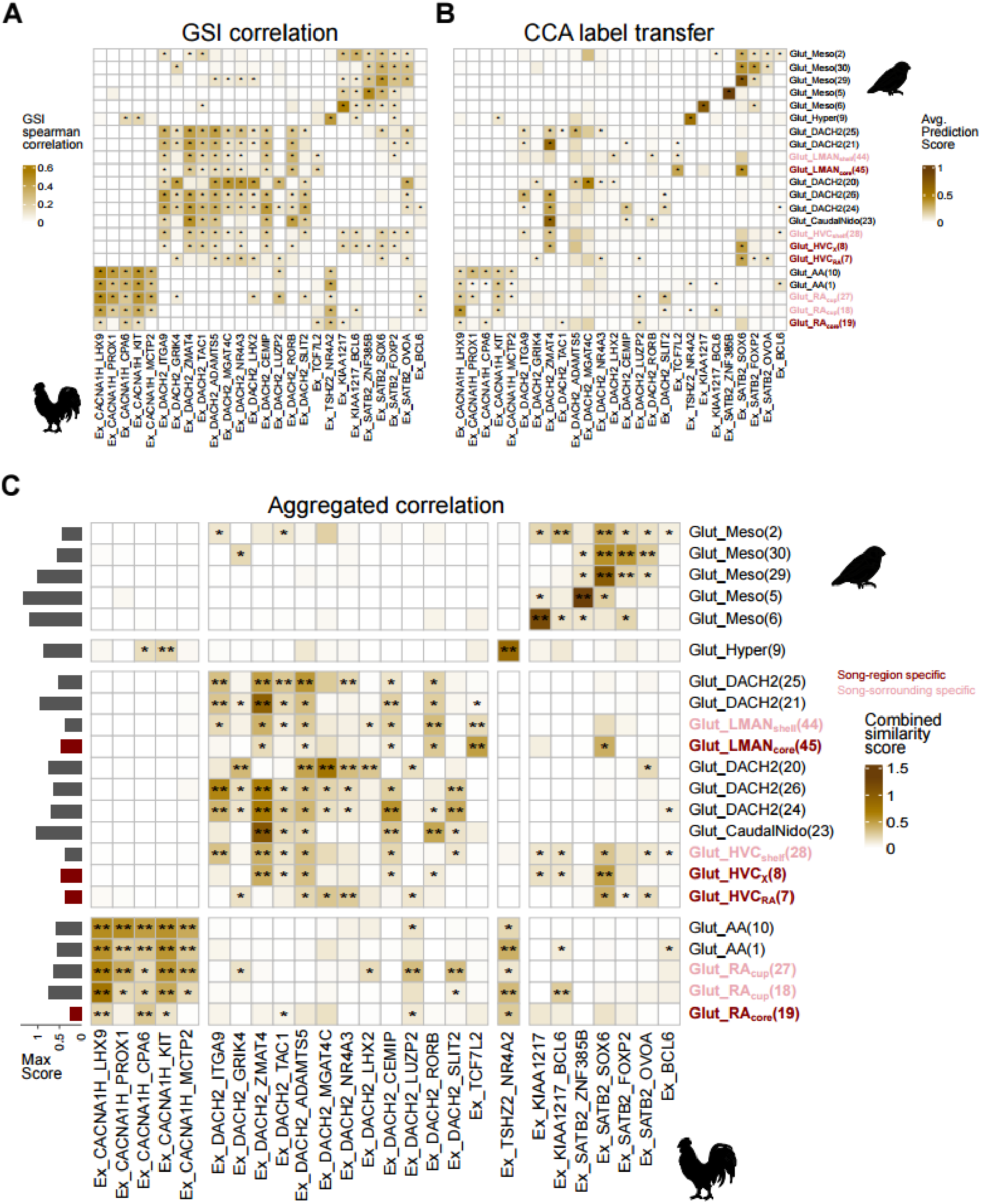
Cluster-level comparison of glutamatergic neurons between chicken and zebra finch pallium. Comparison between glutamatergic clusters in the chicken pallium to clusters of glutamatergic neurons in the zebra finch pallium based on either (**A**) gene specificity index, Spearman correlation, or (**B**) label transfer average prediction score method. Scores in (A) and (B) are aggregated into a single combined score in (**C**).

**Extended Data Fig. 6.**
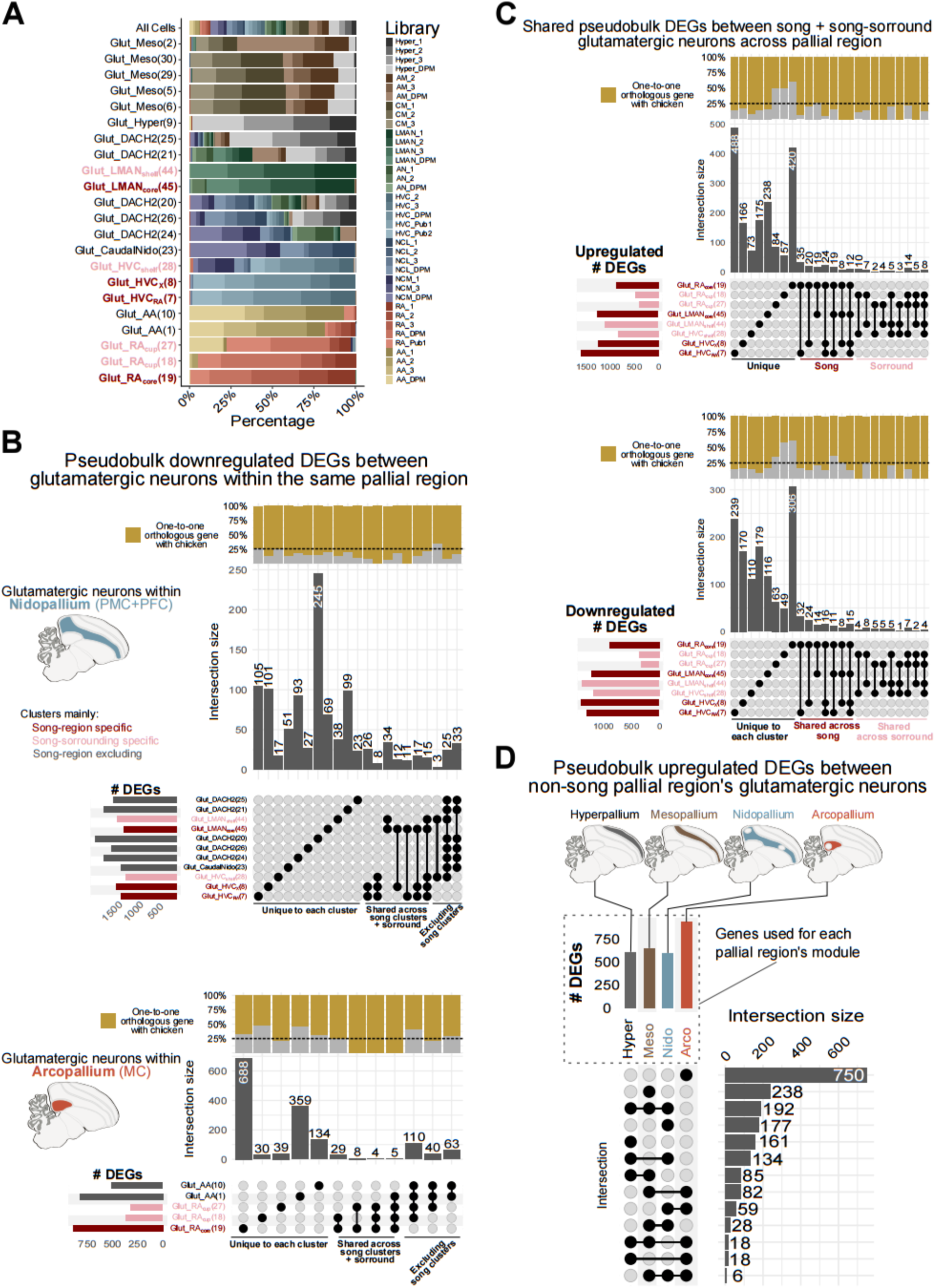
Differential gene expression among glutamatergic neurons of the zebra finch pallium. (A) Stack plots showing the proportion of cells profiled by library per glutamatergic neuron cluster. (B) Upset plot of marker genes for nidopallium glutamatergic neuron clusters compared with other nidopallium glutamatergic neurons (top) and for arcopallium glutamatergic neurons (bottom). The top vertical bars show the percentage of genes with 1-to-1 orthologs in the chicken genome in the corresponding intersection. (C) Upset plot of marker genes of glutamatergic neuron clusters enriched in either vocal imitation regions or shell areas. The top vertical bars show the percentage of genes with 1-to-1 orthologs in the chicken genome in the corresponding intersection. Top, upregulated, and bottom, downregulated. (D) Upset plot of differentially expressed genes between glutamatergic neurons of the four pallial regions of the zebra finch, excluding the vocal imitation regions; used for pallial region module scoring.

**Extended Data Fig. 7.**
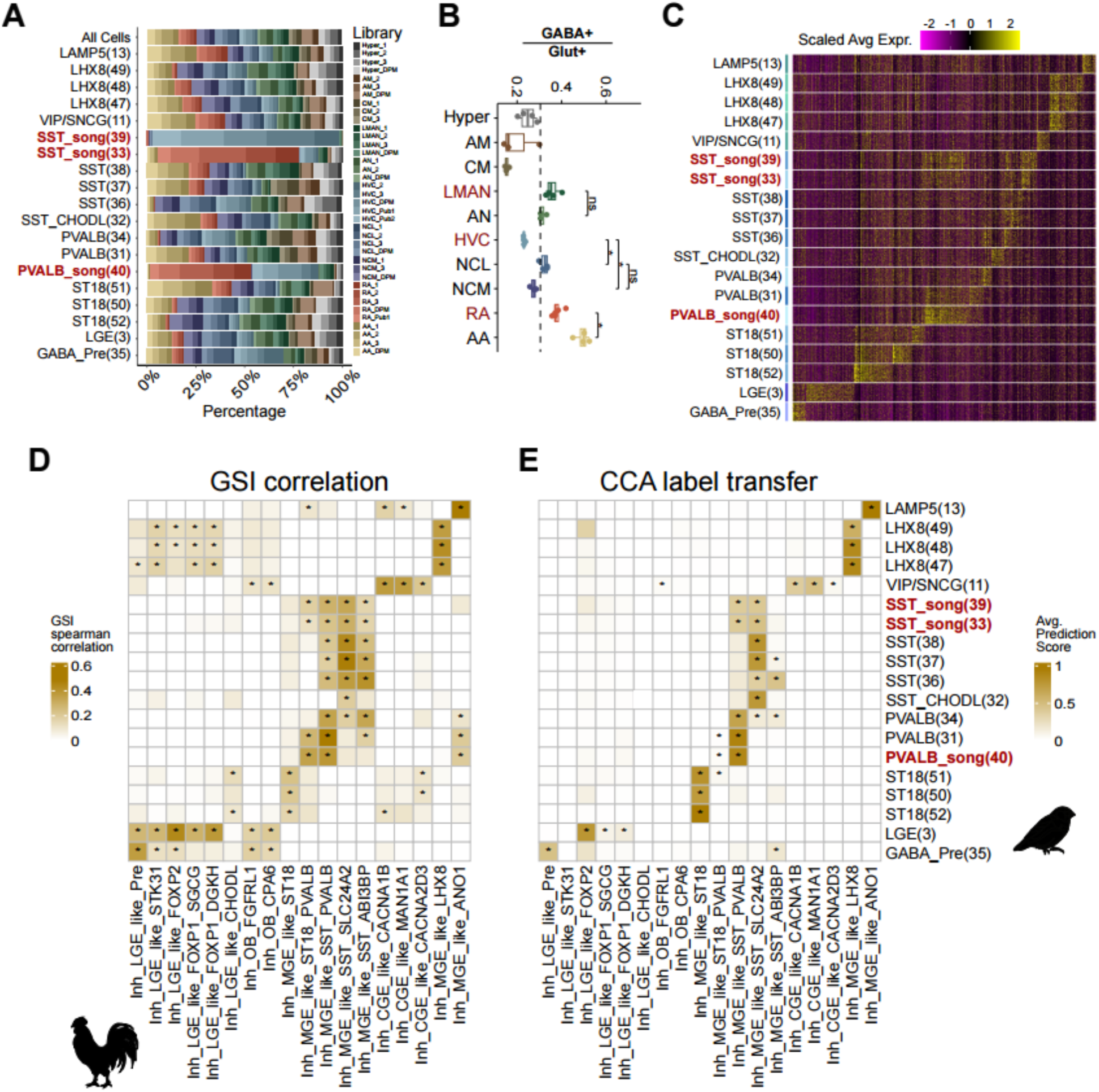
Comparison of GABAergic neurons between chicken and zebra finch pallium. (A) Stack plots showing the proportion of cells profiled by library per GABAergic neuron cluster. (B) Ratio of GABAergic to glutamatergic neurons shown by region, with Wilcoxon tests comparing song regions to their anatomical counterparts. Dashed line indicates the average ratio across all regions. (C) Heatmap of scaled average log-expression of differentially expressed upregulated genes. Sampled from 200 neurons per cluster. Color lines to the left corresponds to assigned cluster color. (**D-E**) Comparison between GABAergic supertypes in the chicken pallium to clusters of GABAergic neurons in the zebra finch pallium based on either (A) gene specificity index, Spearman correlation or (B) label transfer average prediction score method.

**Extended Data Fig. 8.**
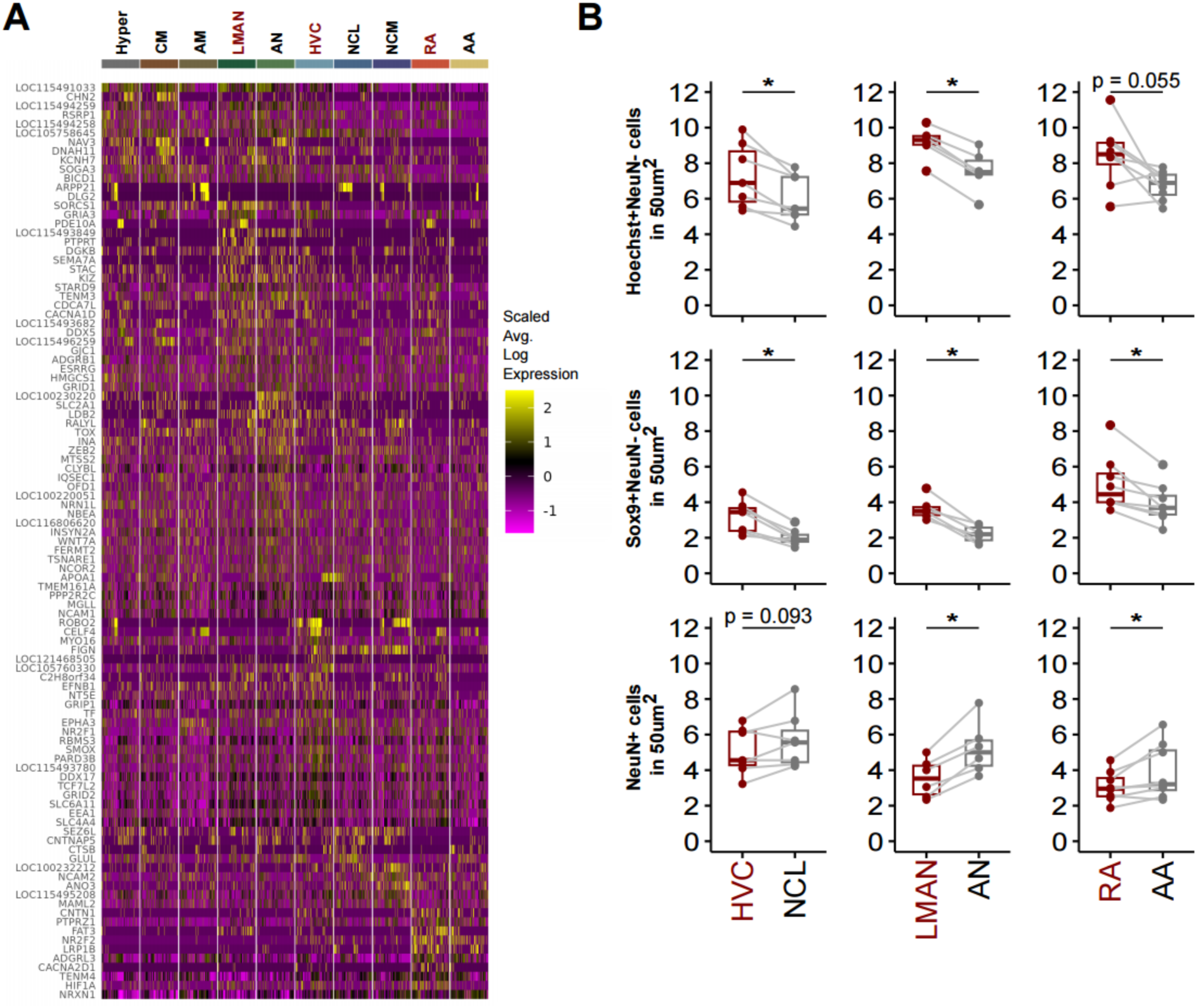
Astrocyte snRNA-seq DEG expression by region and raw IF counts. (A) Heatmap of astrocyte differentially expressed genes by region. (B) Average number of cells by region per 50um^2^ area using IF to identify glia (top), astrocytes (middle), or neurons (bottom).

**Supplementary Table 1**

Sample information and demographics. This table summarizes information per sample tissue, including demographics and sequencing information. It contains information for snRNA-seq and sptRNA-seq

**Supplementary Table 2**

Inter-region glutamatergic cluster composition comparison. Percentages of nuclei in each cluster, divided by region or library.

**Supplementary Table 3**

Glutamatergic neurons differentially expressed genes. Includes pairwise comparisons of glutamatergic clusters within each pallial region and the derived tables used for upset plots. Statistics were obtained using DESeq2 on the aggregated count matrix (Methods).

**Supplementary Table 4**

Pallial region expression modules. This table reports differentially expressed genes of glutamatergic neurons per pallial region, excluding song regions, from the snRNA-seq experiments. Statistics were obtained using DESeq2 on the aggregated count matrix (Methods). It also includes the table used for upset plots. These genes were also used for module scoring in Figure 3C.

**Supplementary Table 5**

GABAergic cluster compositions. Percentages of cells/nuclei in each cluster, divided by region or library. Also included are ratio comparisons and statistics of song-specific clusters to all GABAergic neurons.

**Supplementary Table 6.**

GABAergic differentially expressed genes. Wilcoxon rank-sum differential expression analysis using FindAllMarkers (logfc.threshold = 0.1, min.pct = 0.05, min.diff.pct = 0.15).

**Supplementary Table 7**

DEGs between song-specific, zebra finch widespread, and chicken PVALB and SST GABAergic neurons. Statistics were obtained using DESeq2 on the aggregated count matrix (Methods).

**Supplementary Table 8.**

Glia and astrocyte compositions. Cell counts, percentages, ratios, and statistics for comparisons of glia and astrocytes in song regions and in the chicken for both snRNA-seq and IF. Also included are zebra finch astrocyte inter-region differentially expressed genes. Wilcoxon rank-sum differential expression analysis using FindAllMarkers (logfc.threshold = 0.1, min.pct = 0.05, min.diff.pct = 0.15).

## Notes

https://github.com/cgorozco/Songbird_Pallium_snRNA_sptRNA

### Competing Interest Statement

The authors have declared no competing interest.

